# Parasitic success and venom composition evolve upon specialization of parasitoid wasps to different host species

**DOI:** 10.1101/2020.10.24.353417

**Authors:** Fanny Cavigliasso, Hugo Mathé-Hubert, Jean-Luc Gatti, Dominique Colinet, Marylène Poirié

**Author notes:** these authors should be considered joint senior author (Dominique Colinet, Marylène Poirié).

## Abstract

Female endoparasitoid wasps usually inject venom into hosts to suppress their immune response and ensure offspring development. However, the parasitoid’s ability to evolve towards increased success on a given host simultaneously with the evolution of the composition of its venom has never been demonstrated. Here, we designed an experimental evolution to address this question. We crossed two parasitoid lines of *Leptopilina boulardi* differing both in parasitic success on different *Drosophila* hosts and venom composition. F_2_ descendants were reared on three different *Drosophila* species for nine generations. We tested for evolution of parasitic success over the generations and for the capacity of parasitoids selected on a given host to succeed on another host. We also tested whether the venom composition - based on a statistical analysis of the variation in intensity of the venom protein bands on SDS-PAGE 1D - evolved in response to different host species. Results showed a specialization of the parasitoids on their selection host and a rapid and differential evolution of the venom composition according to the host. Overall, data suggest a high potential for parasitoids to adapt to a new host, which may have important consequences in the field as well in the context of biological control.

## 1. Introduction

Endoparasitoid wasps are insects whose larvae develop inside the host, generally leading to its death [1]. High selection pressures are therefore exerted to maximize the parasitic success that has been shown to evolve rapidly according to host resistance [2–4] and host species [5,6]. The ability to succeed in multiple host species and adapt to a new host is important for the abundance and survival of parasitoids in the event of environmental changes, such as the local extinction of a host. The host range of parasitoids is generally not limited to a single species, even for those considered specialists. For example, *Leptopilina boulardi*, considered a specialist of *D. melanogaster* and *D. simulans*, can develop in other species of the melanogaster subgroup of Drosophilidae, including *D. yakuba* [7,8].

Parasitoids have evolved different strategies to circumvent the host’s immune defense, which usually consists of the formation of a multicellular melanized capsule around the parasitoid egg, together with the activation of the phenoloxidase cascade which leads to the production of melanin and the release of toxic radicals [9–11]. The most common strategy is the injection of venom with the egg, which suppresses the encapsulation process [12–14]. To our knowledge, only two studies have analyzed the ability of parasitoid venom to evolve according to the host. Both *Lysiphlebus fabarum* parasitic success and expression of venom genes rapidly and differentially evolved depending on the strains of a defensive symbiont hosted by the host *Aphis fabae* [4]. More recently, a rapid and differential evolution of the venom protein composition of *L. boulardi* was described according to the susceptible/resistant phenotype of two *D. melanogaster* host strains [15]. However, the relationship to parasitic success was not analyzed. Since these studies involved a single host species, our goal here was to investigate whether the parasitic success and the venom protein composition could evolve according to different host species. We also sought to determine whether the venom of a parasitoid wasp successful in several host species contains broad-spectrum factors or a combination of specialized factors, each specific to a given host.

We used *L. boulardi* as a model because of its intraspecific variability in both venom composition and parasitic success on different hosts. The ISm line always succeeds on *D. melanogaster* but is consistently encapsulated by *D. yakuba* whereas the ISy line can succeed on both Drosophila species but only on certain genotypes [16]. The venom of these lines differs widely, mainly due to quantitative differences in the venomous proteins. As an example, a RhoGAP domain-containing protein named LbGAP is in a significantly higher amount in the venom of ISm than in that of ISy [17,18]. LbGAP would be necessary for ISm parasitic success on an ISy-resistant *D. melanogaster* strain through the induction of morphological changes in the lamellocytes, host immune cells devoted to the encapsulation [17,19–22]. LbSPN, a serine protease inhibitor of the serpin superfamily [18] illustrates the qualitative variation of venom proteins between ISm and ISy. LbSPNy, which inhibits the activation of the phenoloxidase cascade in *D. yakuba* [23] and LbSPNm are one of the most abundant proteins in the ISy and ISm venom, respectively. Although they are encoded by allelic forms of the same gene, they differ in molecular weight and the sequence of the active site, suggesting they have different targets and/or function [18].

Here, we report data from an experimental evolution designed to evaluate the evolvability of *L. boulardi* (i) parasitic success and (ii) the venom protein composition according to different *Drosophila* host strains and species. The experiment was designed to characterize the venom allowing the parasitoid to develop on different hosts and to inform on whether such venom contains rather broad-spectrum factors or a combination of factors specific to each host. For this, we crossed *L. boulardi* ISm and ISy individuals and reared their descendants for nine generations independently on the hosts species and strains differing for their resistance to these wasps. We then analyzed the parasitic success on the selection host and the venom composition in three generations: after the first (F_3_) and the last (F_11_) generation under selection and after an intermediate generation (F_7_). We also tested for a specialization of parasitoids by measuring the parasitic success of individuals from the latest generation of selection on all the different hosts. Overall, parasitoids have shown specialization to their selection host as well as a rapid and differential evolution of the venom composition. Changes in the intensity of the venom protein bands were mainly observed in response to selection on a single host, although some changes occurred under selection by several hosts. This suggests that most of the venom factors of this wasp are host-specific while a few may have a wider spectrum.

## 2. Methods

### 2.1. Biological material

*L. boulardi* ISm (Gif stock number 431) and ISy (Gif stock number 486) isofemale lines originate from populations collected in Nasrallah (Tunisia) and Brazzaville (Congo), respectively [24]. Both lines were reared on their susceptible *Drosophila melanogaster* maintenance strain (Nasrallah from Tunisia, Gif stock number 1333, here called S_Nasr_) at 25°C. Emerged adults were kept at 20°C on agar medium and fed with honey.

Five *Drosophila* host strains from three different species differing in their resistant/susceptible phenotype to ISm and ISy were used (Figure 1A). The *D. melanogaster* R strain (Gif stock number 1088), originating from isofemale lines obtained from a population of Brazzaville, Congo [25] through subsequent genetic approaches [26,27], is resistant to ISy and susceptible to ISm [28,29]. The *D. simulans* Japan strain, from a population collected in Japan, is susceptible to ISm and ISy. *D. yakuba* 1907 (Gif stock number 1907) and 307 (Gif stock number 307.14), originate from Tanzania and from the São Tomé island, respectively. *D. yakuba* 1907 is resistant to both parasitoid lines while 307 is resistant to ISm and susceptible to ISy.

**Figure 1.**
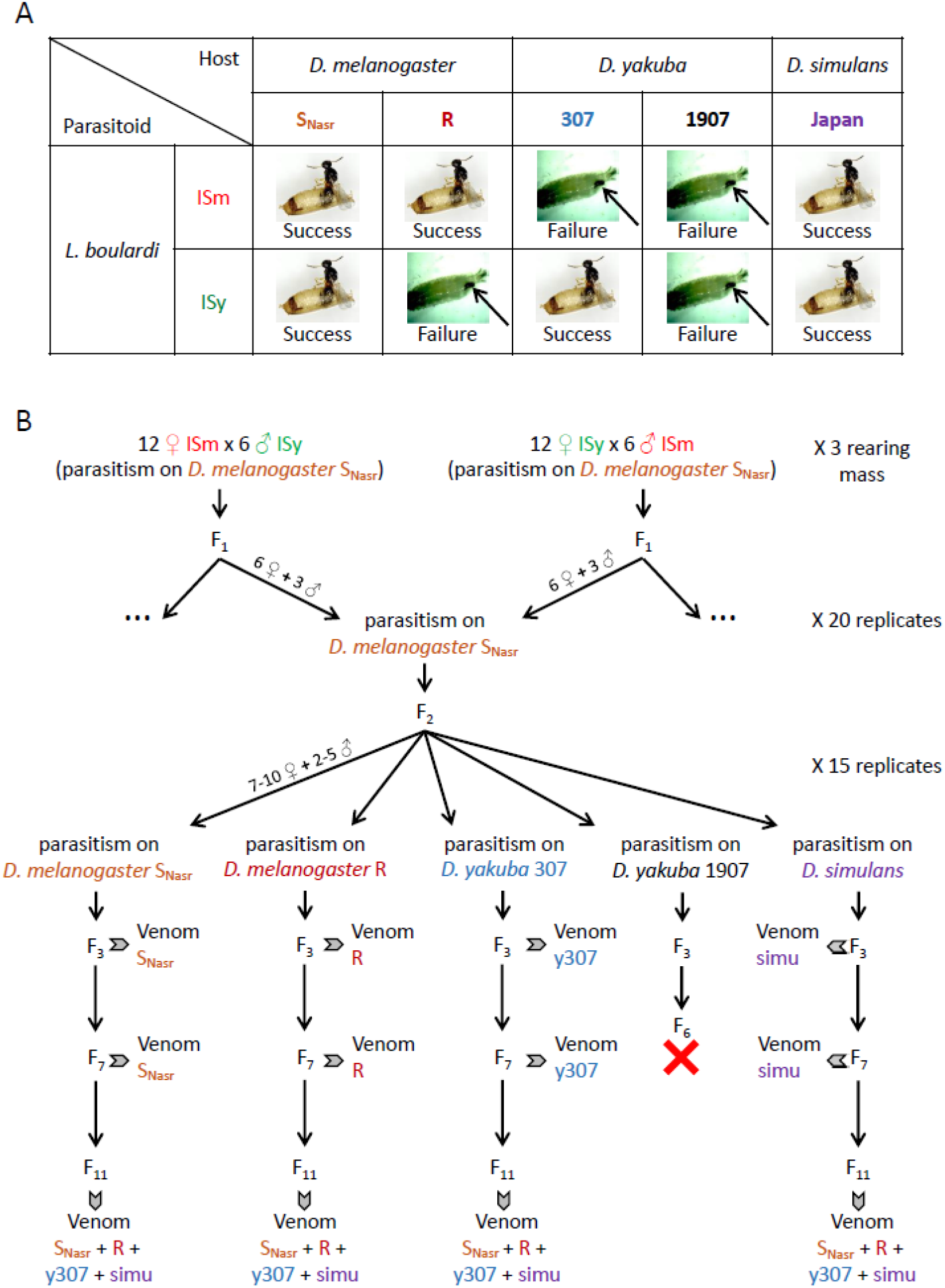
Biological model and experimental evolution protocol. A. Outcome of interaction between the five host strains (*D. melanogaster* SNasr, *D. melanogaster* R, *D. yakuba* 307, *D. yakuba* 1907 and *D. simulans*) and the two *L. boulardi* lines (ISm and ISy). The black arrow shows an encapsulated parasitoid egg inside a *Drosophila* larva. B. Design of the experimental evolution. ISm and ISy: ISm and ISy lines of *L. boulardi*; SNasr: *D. melanogaster* SNasr; R: *D. melanogaster* R; y307: *D. yakuba* 307; simu: *D. simulans*. The red cross indicates the extinction of parasitoids reared on *D. yakuba* 1907, therefore no analysis was performed on individuals from this host. F3, F7 and F11: the three generations of *L. boulardi* that were analyzed for venom composition and parasitic success. The F3 is the first generation under selection.

### 2.2. Experimental evolution protocol

Three wasp mass rearings were created from (♀ISm x ♂ISy and ♀ISy x ♂ISm) crosses of 12 virgin females and six males on the *D. melanogaster* S_Nasr_ host (Figure 1B). A total of 15 replicates were then created from these mass rearings using six F_1_ hybrid females and three F_1_ males from each direction of crossing, still on *D. melanogaster* S_Nasr_. Finally, F_2_ individuals from each of the 15 replicates were used to produce five groups that were then reared separately on the five different hosts until the F_11_ generation (F_6_ only for *D. yakuba* 1907, therefore excluded for further analyses; Figure 1B). For each of the 75 populations (15 replicates x five hosts), up to 10 females and five males (mean: 9.51 ♀, 4.31 ♂) were randomly chosen to produce the F_3_ generation, and up to 20 females and 10 males (mean: 18.48 ♀, 8.99 ♂) to produce F_4_ to F_11_ generations.

The parasitic success and venom composition were analyzed for females of three generations: the first generation of selection on different hosts (F_3_), an intermediate generation (F_7_), and the last generation (F_11_). Since the venom composition and the parasitoid behavior could vary between females depending of the number of eggs previously laid, only females never allowed to oviposit were used for the analyses.

### 2.3. Analysis of the outcome of the *Drosophila – L. boulardi* interaction

#### Parasitic tests and dissection

Twenty second-instar larvae of the investigated host species were deposited in small medium-containing dishes with one parasitoid female. The parasitoid was removed after 4h at 25°C and the dishes kept at 25°C for 48h until dissection of the *Drosophila* larvae. They were then categorized as (i) non-parasitized, (ii) mono-parasitized and (iii) multi-parasitized (see Figure S1 for proportions of each). Only mono-parasitized host larvae were considered for the analysis to avoid unpredictable effects of the presence of several parasitoid larvae in a single host.

Three possible outcomes were then recorded: i) free parasitoid larva alone, ii) free parasitoid larva together with an open capsule and iii) complete closed capsule (Figure 2A). Among these, the percentage of outcomes for i) and iii) were generally used to assess the parasitoid’s immune suppression ability and the host encapsulation capacity [5,25,30]. In this work, we added the outcome ii) to evaluate the parasitoid’s ability to escape the encapsulation process initiated by the host after recognition of the parasitoid egg. Since the escaped parasitoid larva was alive, we considered this outcome to be a parasitic success, similar to a free parasitoid.

**Figure 2.**
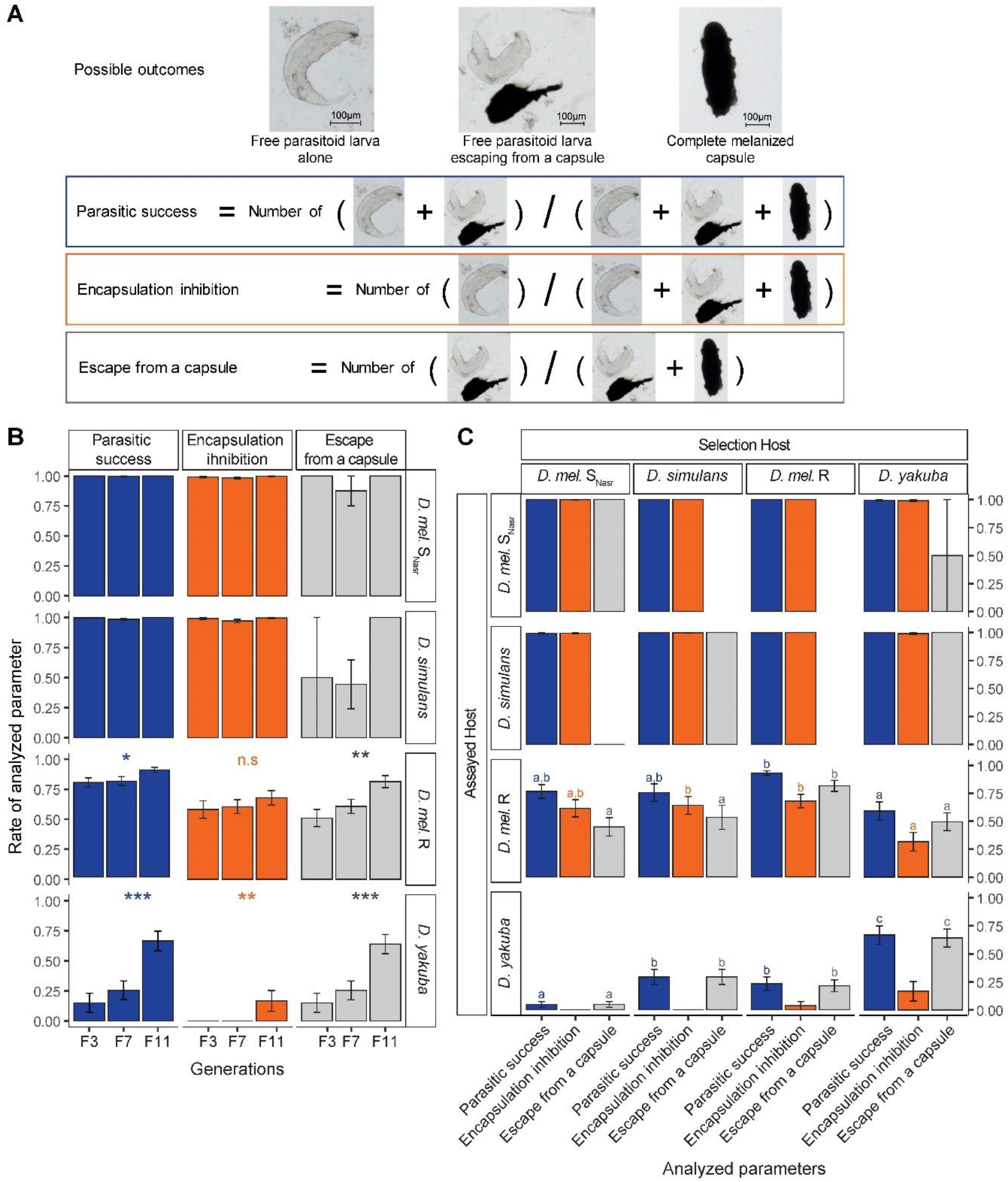
Evolution of the parasitoid’s ability to bypass the encapsulation response of the host. A. Possible outcomes observed in *Drosophila* larvae 48 hours after parasitism and details of the analyzed parameters: parasitic success, parasitoid capacity to inhibit encapsulation and parasitoid capacity to escape from a capsule. B. Evolution of (i) parasitic success, (ii) parasitoid capacity to inhibit the egg encapsulation by the host or (iii) parasitoid capacity to escape from a capsule, depending of the selection host. C. Capacity of parasitoids from the F11 generation to bypass encapsulation by four different hosts. The hosts listed on the top are the “selection hosts”, those listed on the left are the “assayed hosts”, used for parasitism assays. *D. mel*.: *D. melanogaster*. Letters above bars indicates the significance of the difference. Different color of letters indicate different statistical tests. Error bars indicate standard errors.

#### Statistical analysis

We used a generalized linear mixed model (GLMM) to analyze each parameter with the continuous generation or the selection host as a fixed effect, the replicates as a random effect and a binomial error distribution. The GLMMs were fitted with the *glmer* function in “lme4” R package [31], except for unbalanced data for which they were fitted with the *bglmer* function implemented in the “blme” R package [32]. As for all the binomial GLMMs performed, overdispersion was tested with the *overdisp_fun* function [33] and accounted for when necessary (p < 0.05) by adding a random factor corresponding to the observations number [34]. GLMMs were followed by post-hoc Tukey tests with the “multcomp” R package [35] to compare hosts two by two.

### 2.4. Venom separation, western blot and data acquisition

*L. boulardi* venom reservoirs were dissected individually in 15 µl of insect Ringer supplemented with protease inhibitors cocktail (PI; Roche), mixed with an equivalent volume of Laemmli reducing buffer and heated (95°C, 10 min). The individual protein samples were then split in two, half being used for the global analysis, half for the specific analysis (see the two next paragraphs). Protein separation was done by 1D SDS-PAGE electrophoresis using commercial gels to ensure reproducibility (8–16% Criterion™ TGX™ Precast Midi Protein Gel, Bio-Rad). A sample of venom of ISm and ISy lines, equivalent to half a female reservoir (from a pool of venom from 60 individuals collected in 1.8 ml) was also loaded on each gel and served as a reference. The global analysis was performed as described in [36] and used in [15]. Briefly, the gels were silver-stained and their digital pictures analyzed with the Phoretix 1D software (now CLIQS, TotalLab, UK) to extract the intensity profile of each lane (individual sample). We then used R functions to obtain for each lane the intensities of a set of “reference bands” of known molecular weight. These intensities were normalized and estimated with the following combination of parameters: without background – peak height – quantile normalization (more details in [36]). The normalized intensities of the reference bands are the variables characterizing the venom composition.

The specific analysis was performed using Western blots and antibodies targeting previously characterized *L. boulardi* venom proteins as described in [15]. The LbSPN, LbGAP and LbGAP2 polyclonal antibodies are described in [19,20,37,38]. The secondary was a goat anti-rabbit IgG horseradish peroxidase conjugate (1:10,000; Sigma). Western blots were revealed with a luminescent substrate (Luminata Crescendo; Millipore) and digitalized. Data were then recorded as follows for each individual:

i. LbSPN: the genotype at the *lbspn* locus was determined by the presence/absence of the LbSPNy (54 kDa) and LbSPNm (45 kDa) bands.
ii. LbGAP: a strong signal was observed in the venom of ISm or (ISm × ISy) F_1_ only. The presence/absence of a signal at the expected size was then used to distinguish *lbgap* homozygotes and *lbgap*/*lbgap_y_* heterozygotes from *lbgap_y_* homozygotes.
iii. LbGAP2: the variation in quantity was more continuous. The normalized quantity of LbGAP2 was estimated as the ratio between the signal intensity on the Western blot without background and the sum of the intensities of the reference bands (used as a proxy for the amount of protein in the venom samples) in the corresponding lane on silver stained gels.

### 2.5. Statistical analysis for the global analysis of the venom

#### Evolution of the venom composition

We analyzed the evolution of the venom composition using PERMANOVAs (permutational MANOVAs; “vegan” R package; function *adonis2*; [39,40]). PERMANOVAs measure the association between the multidimensional variation of some explained variables (venom) and some explanatory variables, but at the difference of MANOVAs, they don’t compare the correlation structure among different groups. For this analysis, we measured the multidimensional variation through the Euclidean distance and we tested the significance of explanatory variables’ marginal effects with 5,000 permutations. To determine whether the venom composition has evolved on each host separately, PERMANOVAs were fitted with the generation (F_3_, F_7_ or F_11_; continuous variable) and the population (same as replicates in this situation) as explanatory variables. For *D. yakuba* 307, we fitted an additional PERMANOVA without F_3_ individuals due to their small number. We then determined whether the venom composition has evolved differently depending on the host by fitting a PERMANOVA with the following explanatory variables: the generation, the host, the interaction of both and the 44 experimental populations (sum of replicates maintained on each host until the end of the experiment) to account for the effect of genetic drift. Then, PERMANOVAs were performed to compare the venom composition two by two between hosts to determine whether a venom composition evolved on one host differed from that evolved on another.

To characterize the evolutions detected by the PERMANOVAs, we used linear discriminant analyses (LDA). Specifically, we performed LDA to characterize (i) the evolution of the venom composition on each host separately (LDAs performed with the three groups of individuals i.e. from the three analyzed generations) and (ii) the differential evolution of venom composition depending on the host (LDAs performed with the 12 groups of individuals (three generations × four hosts) and each two by two comparison of venom composition between hosts. We used the “ade4” R package [41] with the individual venom compositions as continuous variables and the generation or the “host × generation” interaction as a factor. Since LDA does not account for the variation between replicates, they were centered before the analysis using the *wca* function (within class analysis) in the “ade4” R package. With this additional step, all replicates had the same mean for each variable and therefore the variation among generations and hosts cannot result from the variation from replicates and unbalanced data. The LDA axes were labeled with the biological meaning determined from the position of the 12 groups of individuals (generations × hosts) on these axes. To describe the venom evolution trend, we plotted for each host in each LDA an arrow representing the linear regression fitted to coordinates of the three centroids corresponding to the F_3_, F_7_ and F_11_ generations and weighted by the number of individuals per generation and host. The only exception was the LDA on *D. yakuba* 307 for which the linear regression was calculated from centroid points of the F_7_ and F_11_ generations only.

#### Evolution of venom protein bands

Non-parametric Spearman rank correlation tests were performed for each LDA done for each host separately to identify the protein bands that correlated with the regression describing the trend of venom evolution. P-values were Bonferroni corrected using the *p.adjust* R function.

As some protein bands were probably indirectly selected due to their correlation with other bands, a combination of clustering and partial correlation analyses was used to identify bands undergoing direct selection. We first performed an UPGMA clustering analysis (“hclust” R package) with 1 - |ρ| as the metric distance, where ρ is the spearman correlation. Then, we used a threshold correlation of 0.4 to construct band clusters for which false detection of certain bands as “correlated” could have occurred. For each cluster with at least two protein bands correlated to the regression (represented by an arrow), we performed partial correlations with the *pcor* function in the “ppcor” R package to determine if the residual variation in the intensity of each band, independent from the other bands of the cluster, was still correlated to the regression describing the evolution. P-values were also Bonferroni corrected using the *p.adjust* R function.

#### Comparison of the venom composition evolved on each host to that of the parental lines

To make this comparison, we computed the averages of venom composition evolved on each host separately for each replicate. Each of the 44 populations from the first and last generation of selection (F_3_ and F_11_) was scaled between 0 and 1 depending on its distance to the ISm or ISy venom using the formula *D_ISy_* /(*D_ISm_* + *D_ISy_*), *D_ISy_* and *D_ISm_* being the Euclidean distances between the average venom composition of the population and the venom composition of ISm and ISy. Then, we tested whether the venom composition of parasitoids reared on each host evolved toward a parental line by comparing the scaled distances between F_3_ and F_11_ generations with paired Wilcoxon rank tests. Finally, to determine whether a protein band selected on a given host corresponded to an ISm or ISy band, we assigned it a value from 0 to 1 in relation to a higher intensity in ISy or ISm, respectively. This value is the intensity of the band in the venom of ISm divided by the sum of its intensities in the venom of ISm and ISy.

### 2.6. Statistical analysis for the specific analysis of venom

The three variables describing LbSPN, LbGAP and LbGAP2 are of different type (categorical for LbSPN with two different alleles, presence/absence for LbGAP, continuous for LbGAP2 with the relative intensity). We therefore used different approaches to analyze them.

LbSPN is a codominant marker with two alleles (*lbspnm* and *lbspny*) encoding proteins of distinct molecular weight. To determine if the frequency of these alleles had evolved on each host, we fitted one generalized linear mixed model (GLMM) per host with generation as a fixed continuous effect and replicates as a random effect with a binomial distribution using the “lme4” R package [31].

LbGAP is a dominant marker with two alleles (*lbgap* and *lbgap_y_*) determining the presence in the venom of the LbGAP protein in detectable quantity (*lbgap* homozygotes and *lbgap*/*lbgapy* heterozygotes) or not (*lbgapy* homozygotes). The evolution of the presence/absence of the LbGAP/LbGAPy proteins independently on each host was tested using the same procedure as for LbSPN.

For LbGAP2 (continuous variation), we used linear mixed models (LMM) with generation as a fixed continuous effect and replicates as a random effect to determine if the quantity of LbGAP2 had evolved, separately for each host. The models were fitted with the “nlme” R package [42] on the box cox-transformed (lambda = 0.25) standardized intensity of LbGAP2 to normalize the residues.

## 3. Results

### 3.1. Experimental evolution protocol

The variability on which the selection took place was generated by reciprocal crosses between the ISm and ISy parasitoid lines. F_1_ hybrids were then used to produce the 15 F_2_ replicates from which five groups were formed and reared independently on *D. melanogaster* S_Nasr_, *D. melanogaster* R, *D. yakuba* 307, *D. yakuba* 1907 and *D. simulans* (Figure 1B). Of these 75 starting F_2_ experimental wasp populations, 44 were successfully maintained until the F_11_ generation. Among these, none of the *D. yakuba* 1907 replicates survived after the F_6_, preventing further analyses for this host. 12 and 15 replicates survived on *D. melanogaster* S_Nasr_ and *D. simulans*, respectively (Table S1), and 13 on *D. melanogaster* R, whereas only four could be maintained on *D. yakuba* 307 until the F_11_, the other replicates mainly becoming extinct at the first generation of selection (F_3_) (Table S1).

### 3.2. Evolution of the interaction outcome according to the selection host

We tested the evolution of the parasitoids capacity to bypass the encapsulation response of their selection host after five and nine generations of selection (i.e. F_7_ and F_11_), using two females per experimental population, except for *D. yakuba* 307 for which four females were tested since fewer replicates were available. Three parameters were analyzed: (i) the parasitic success, i.e. among mono-parasitized host the proportion of hosts containing a free parasitoid larva alone or together with an open capsule, (ii) the capacity of the parasitoid to inhibit the capsule formation, i.e. among mono-parasitized host the proportion of *Drosophila* larvae containing a free parasitoid larva alone and (iii) the capacity of the parasitoid to escape from the capsule, i.e. among mono-parasitized hosts showing an encapsulation response, the proportion of hosts containing a free parasitoid larva together with an open capsule (Figure 2A).

The parasitic success of parasitoids reared on *D. melanogaster* S_Nasr_ or *D. simulans* on this same host remained close to 100% throughout the experimental evolution (Figure 2B, Table S2, GLMM, p > 0.30 for both hosts). In contrast, it increased with the generation on *D. melanogaster* R (from about 80% at F_3_ to 90% at F_11_) and *D. yakuba* 307 (from about 20% at F_3_ to 65% at F_11_) (Figure 2B, Table S2, GLMM, p = 0.002 for *D. melanogaster* R and p < 0.001 for *D. yakuba* 307). For *D. melanogaster* R this increase seemed to result solely from the increased capacity to escape from the capsule (Figure 2B, Table S2, GLMM, p = 0.001) since no significant change was observed for its ability to inhibit encapsulation (Figure 2B, Table S2, GLMM, p = 0.33). For *D. yakuba* 307, the success increase seemed to result mainly from a higher capacity to escape from the capsule (Figure 2B, Table S2, GLMM, p <0.001) but also, to a lesser extent, to inhibit encapsulation at F_11_ (Figure 2B, Table S2, GLMM, p = 0.003). Finally, a much lower parasitic success was observed at F_3_ for *D. yakuba* 307 (about 20%) than for the other hosts (about 80% to 100% depending on the host) (Figure 2B).

### 3.3. Specialization of parasitoids to their selection host

To determine whether the change in parasitoid ability to bypass host encapsulation was specific to the selection host, we compared the success of F_11_ parasitoids on their own selection host *vs.* the three other hosts. The parasitic success on *D. melanogaster* S_Nasr_ and *D. simulans* was close to 100% regardless of the selection host (Figure 2C, Table S3, GLMM, Tukey post hoc-test, p > 0.70 for both hosts). In contrast, parasitoids reared on *D. melanogaster* R were more successful on this same host than those reared on *D. yakuba* 307 (Figure 2C, Table S3, GLMM, Tukey post hoc-test, p <0.001). This can be explained by a higher capacity of parasitoids selected on *D. melanogaster* R to both inhibit encapsulation and escape from a capsule compared to those selected on *D. yakuba* 307 (Figure 2C, Table S3, GLMM, Tukey post hoc-test, p = 0.015 for encapsulation inhibition and p < 0.001 for escape capacity). Finally, parasitoids selected on *D. melanogaster* R had a higher capacity to escape from a capsule of this same host compared to parasitoids selected on *D. melanogaster* S_Nasr_ or *D. simulans*, despite a similar parasitic success (Figure 2C, Table S3, GLMM, Tukey post hoc-test, p < 0.001for the escape capacity between *D. melanogaster* R and the two other hosts; p = 0.09 and p = 0.30 and for the parasitic success between *D. melanogaster* R vs *D. melanogaster* S_Nasr_ or vs *D. simulans*, respectively).

Parasitoids reared on *D. yakuba* 307 had a higher parasitic success on this same host than parasitoids reared on all the other hosts (Figure 2C, Table S3, GLMM, Tukey post hoc-test, p < 0.005 for all comparisons involving *D. yakuba* 307). The parasitoids capacity to inhibit encapsulation by *D. yakuba* 307 was very low, with no significant difference between the selection hosts (Figure 2C, Table S3, GLMM, Tukey post hoc-test, p > 0.60). Therefore, the difference in parasitic success between selection hosts was most probably explained by an increase capacity to escape from the capsule (Figure 2C, Table S3, GLMM, Tukey post hoc-test, p < 0.05 for each pairwise comparison except for *D. melanogaster* R and *D. simulans* for which p = 0.76).

### 3.4. Differential evolution of venom composition according to the host

The venom analysis was performed on 50 individuals per host and generation distributed over all replicates, except for *D. melanogaster* S_Nasr_ at F_3_ (47 ♀), R at F_11_ (49 ♀) and *D. yakuba* 307 (10, 39 and 40 ♀ at F_3_, F_7_ and F_11_, respectively). In total, the venom protein content was analyzed for 535 females and 36 reference protein bands were identified whose intensities represent the variables describing the venom composition (Figure S2).

The PERMANOVAs performed for each host separately revealed a strong generation effect for *D. melanogaster* S_Nasr_ and R, and *D. simulans* (Table S4, p < 0.01), suggesting that the venom composition evolved in response to each of these hosts. This was confirmed by a linear discriminant analysis (LDA) discriminating the groups of individuals based on generation for the three hosts (Figure 3). Of note, the generation effect revealed by the PERMANOVA was only significant for *D. yakuba* 307 when removing F_3_ individuals from the analysis (Table S4, p = 0.042), likely because only few females were available at F_3_. LDA may overfit groups composed of few individuals [43,44] and was therefore performed for *D. yakuba* 307 only between F_7_ and F_11_. LDA discriminated these two groups, therefore confirming the generation effect revealed by the PERMANOVA (Figure 3).

**Figure 3.**
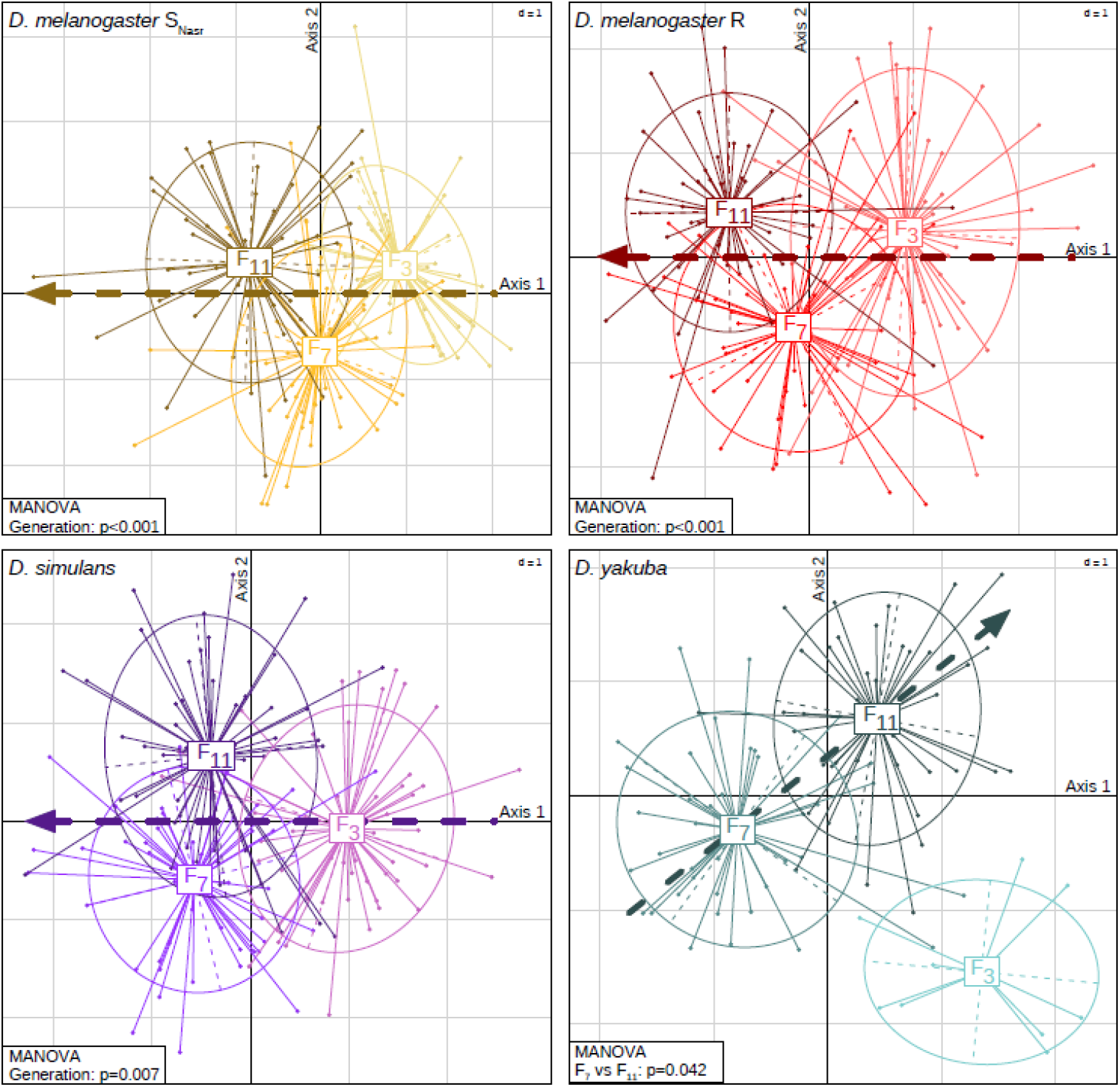
Evolution of the venom composition according to the selection host. Position of the individuals (shown as dots) on the two discriminant axes for each selection host. Individuals are grouped and coloured according to the generation (F3, F7 and F11). The arrows represent the trend of the venom evolution. The dotted arrows represent the linear regressions calculated from coordinates of the three centroid points corresponding to the F3, F7 and F11 generations and weighted by the number of individuals per generation. For *D. yakuba* 307, the linear regression was calculated from centroid points of F7 and F11 only due to the low number of females available in F3. P-values obtained from the PERMANOVA for the “generation” effect are provided at the bottom right of the LDA (see Table S4).

A PERMANOVA fitted to all hosts together evidenced a significant effect of the “host” (p < 0.001), the “generation” (p < 0.001) and their interaction (p < 0.001) on the variation of bands intensity, demonstrating that the venom composition evolved differently on the different hosts (Table S5). Yet, although significant, the combined effects of the “host”, “the generation”, and the interaction of the two only accounted for a small part of the variance of the venom composition (R² = 8.7%; Table S5) compared to that explained by the experimental “population” (R² = 26.6%; Table S5). The LDA confirmed the differential venom evolution according to the host (Figures 4A and 4B, p < 0.001) but the 12 groups (four hosts x three generations) largely overlapped, which confirmed the small part of the variance explained by the host and the generation. The first four out of the 11 discriminant axes were the only ones to be biologically meaningful (Figures 4A and 4B). The first two axes discriminated the F_7_ and F_11_ generation for parasitoids reared on the different hosts, except for the two *D. melanogaster* strains (Figure 4A), evidencing a rapid and differential evolution of the venom composition according to the host species. This was confirmed by the arrows describing the venom evolution that were orthogonal or in opposite direction for the different host species except for the two *D. melanogaster* strains (Figure 4A). Axis 3 discriminated the three generations for parasitoids reared on *D. melanogaster* strains and *D. simulans* while individuals reared on *D. yakuba* 307 were discriminated by axis 4 (Figure 4B). This suggests that the venom of parasitoids reared on *D. yakuba* 307 evolved differently than that of parasitoids reared on the other hosts as confirmed by the direction of the arrow describing the venom evolution for individuals raised on this species (Figure 4B).

**Figure 4.**
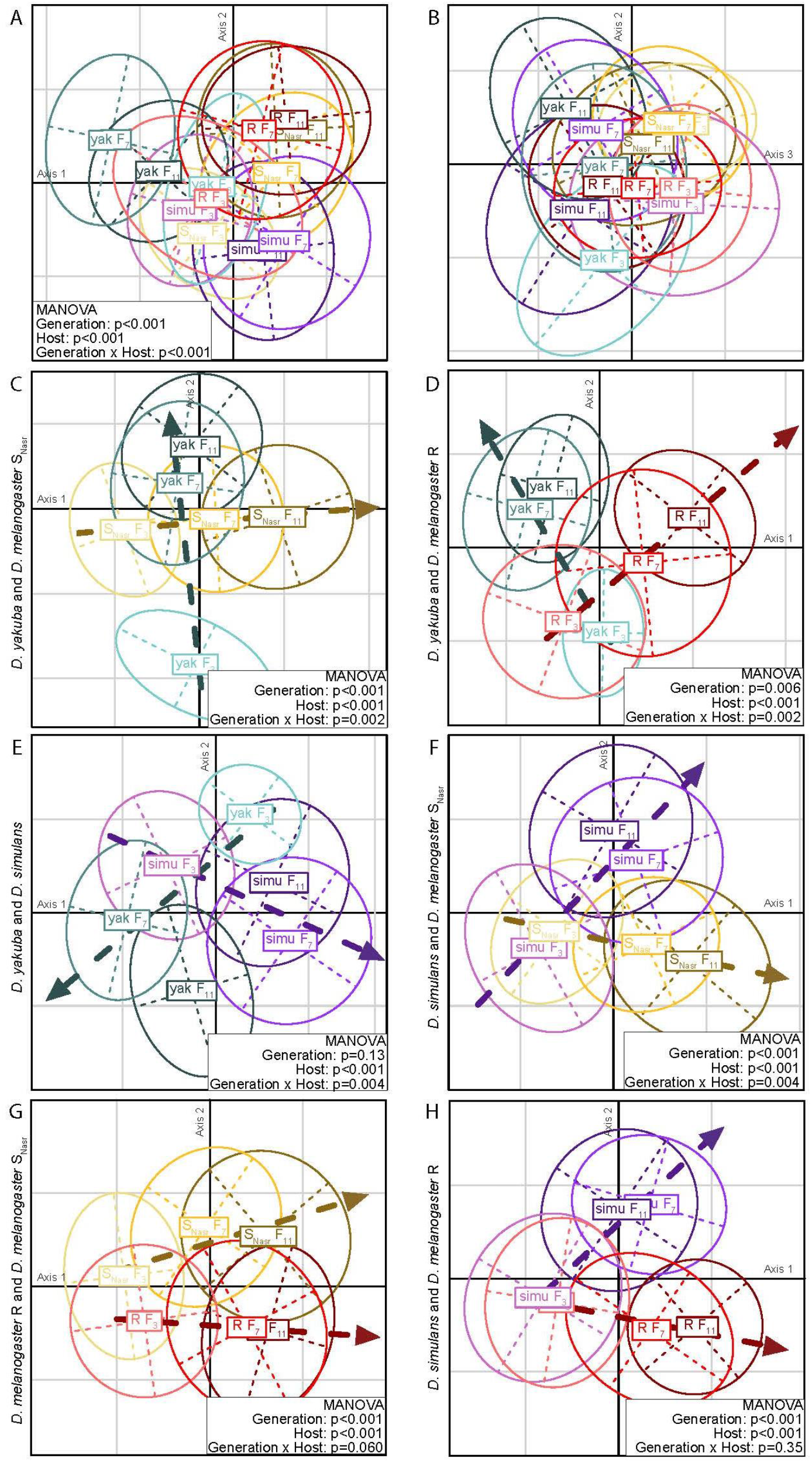
Differential evolution of venom composition. Discriminant analysis showing ellipses and centroids (intersection of dashed lines) of each group formed by the host and the generation are represented (see Figure S4 for more details on the position of each individual). The name of each group is written on the centroid. SNasr: *D. melanogaster* SNasr; simu: *D. simulans*; R: *D. melanogaster* R; yak: *D. yakuba* 307. A-B. Venom evolution in response to the four hosts at F3, F7 and F11 generations on discriminant axes 1 and 2 (A) and discriminant axes 3 and 4 (B). C-H. Discriminant analysis performed on the two first discriminant axes for each two-by-two comparison of venom composition between parasitoids selected on different hosts. For each of these LDAs, the pair of host considered is indicated at the left. Individuals are grouped and coloured according to their selection host and generation. The dotted arrows representing the direction of the venom evolution are the linear regressions fitted to the coordinates of the three centroid points (F3, F7 and F11). P-values obtained from the PERMANOVA for effects of the “generation”, “host” and “generation × host” interaction are provided at the bottom right of the LDA (see Table S5 for more details).

To better identify the hosts on which the venom composition has evolved differentially, PERMANOVAs and LDAs were performed to compare them two by two. There was a strong effect of the “generation × host” interaction for all comparisons involving *D. yakuba* 307 (Table S5, p < 0.01) and the comparison between *D. melanogaster* S_Nasr_ and *D. simulans* (Table S5, p = 0.004). This was confirmed by the LDA since groups of individuals were separated at F_7_ and F_11_ and arrows representing the direction of evolution were almost orthogonal (Figures 4C to 4F). No effect of the “generation × host” interaction was observed in the two PERMANOVAs comparing evolution on *D. melanogaster* R to either *D. melanogaster* S_Nasr_ or *D. simulans* (Table S5, p = 0.06 for the comparison with *D. melanogaster* S_Nasr_; p = 0.35 for the comparison with *D. simulans*). The LDAs nevertheless suggested a trend of differential evolution although the separation of groups at generations F_7_ and F_11_ was lesser between the two strains of *D. melanogaster* than between *D. melanogaster* R and *D. simulans* (Figures 4G and 4H).

### 3.5. Identification of protein bands that evolved on each host

Selected protein bands were identified based on their correlation to the linear regressions (calculated from centroid points of generation groups and represented by arrows) describing the venom evolution in the LDAs performed for each host separately (Figure 3). However, some of these protein bands could have been selected only indirectly because of their correlation with other directly selected bands, either due to their migration proximity on the gel or to a linkage disequilibrium. To disentangle them, we used a combination of clustering (Figure S3) and partial correlation analyses (Table S6) as described in Cavigliasso et al. (2019). The analysis revealed that (i) seven protein bands had evolved on *D. melanogaster* S_Nasr_ (four selected, three counter-selected), (ii) six on *D. melanogaster* R (three selected, three counter-selected), (iii) seven on *D. simulans* (five selected, two counter-selected) and (iv) two on *D. yakuba* 307 (one selected, one counter-selected) (Table 1, Table S6 and Figure S5).

**Table 1.**
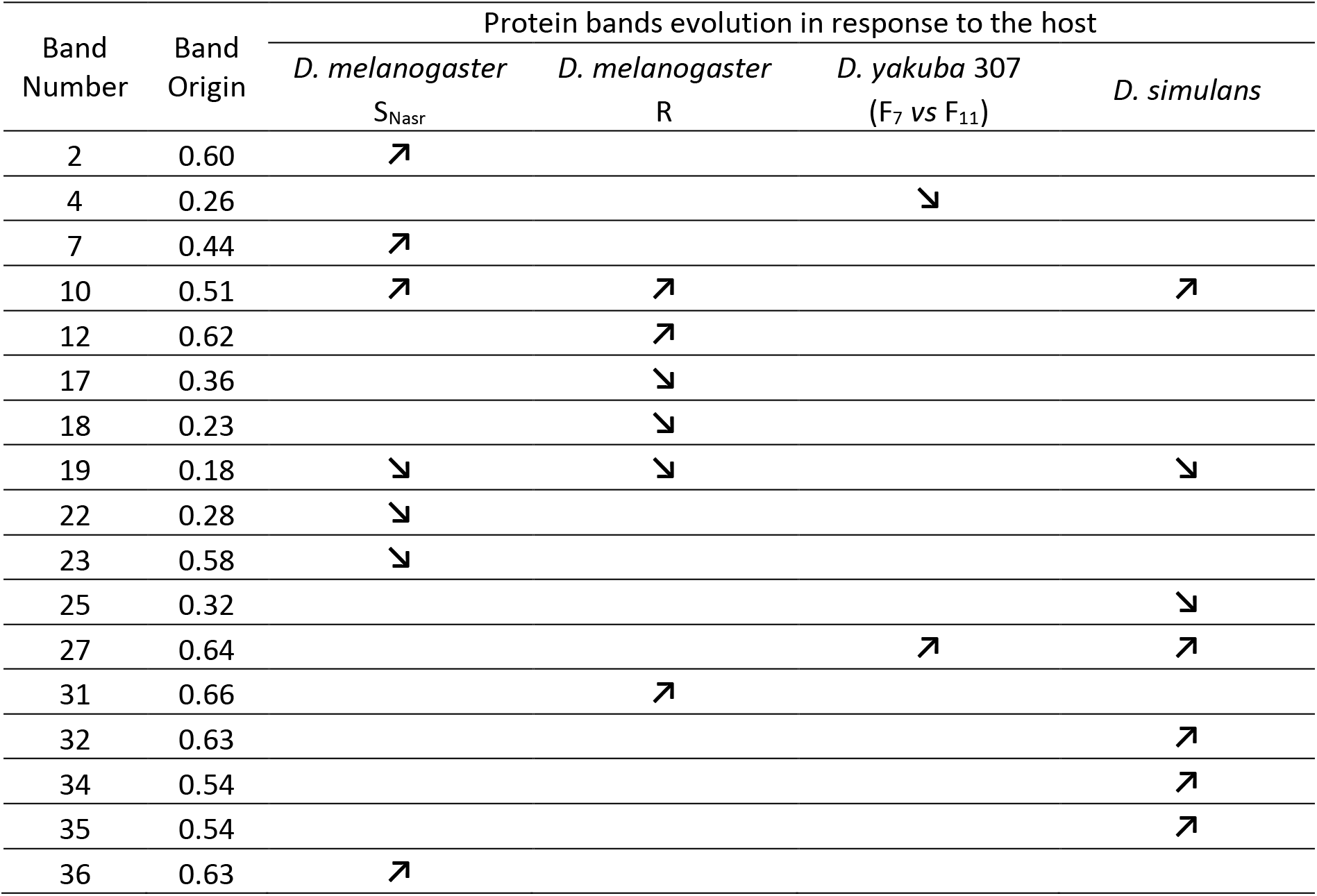
Summary of protein bands evolution in response to the host. Only protein bands correlated to regressions (arrows) in Figure 3 and S5 in response to at least one host are represented. A“↗” corresponds to a selection, a“↘” to a counter-selection in response to the host. The origin of the band was estimated by dividing the band intensity in ISm venom by the sum of its intensity in ISm and ISy venoms; 0 < ISy <0.5; 0.5 < ISm < 1. For the complete data, see Table S6.

| Band Number | Band Origin | Protein bands evolution in response to the host |  |  |  |
| --- | --- | --- | --- | --- | --- |
| | | *D. melanogaster*<br>$S_{Nasr}$ | *D. melanogaster*<br>R | *D. yakuba* 307<br>($F_7$ vs $F_{11}$) | *D. simulans* |
| 2 | 0.60 | ↗ |  |  |  |
| 4 | 0.26 |  |  | ↘ |  |
| 7 | 0.44 | ↗ |  |  |  |
| 10 | 0.51 | ↗ | ↗ |  | ↗ |
| 12 | 0.62 |  | ↗ |  |  |
| 17 | 0.36 |  | ↘ |  |  |
| 18 | 0.23 |  | ↘ |  |  |
| 19 | 0.18 | ↘ | ↘ |  | ↘ |
| 22 | 0.28 | ↘ |  |  |  |
| 23 | 0.58 | ↘ |  |  |  |
| 25 | 0.32 |  |  |  | ↘ |
| 27 | 0.64 |  |  | ↗ | ↗ |
| 31 | 0.66 |  | ↗ |  |  |
| 32 | 0.63 |  |  |  | ↗ |
| 34 | 0.54 |  |  |  | ↗ |
| 35 | 0.54 |  |  |  | ↗ |
| 36 | 0.63 | ↗ |  |  |  |

Of the 17 protein bands that evolved in response to at least one host, only three evolved in the same direction on several hosts. Indeed, bands #10 and #19 were respectively selected and counter-selected on *D. melanogaster* S_Nasr_, *D. melanogaster* R and *D. simulans* (Table 1) and band #27 was selected on *D. simulans* and *D. yakuba* 307. Overall, this confirmed that the venom composition evolved rapidly and differentially between hosts.

### 3.6. Putative identification of proteins that evolved on each host

The tentative identification of the proteins contained in the bands under selection was performed by matching these bands with those in 1D electrophoresis gels used for *L. boulardi* ISm and ISy venom proteomics [18]. The level of intensity of a band can result from that of several proteins having migrated at the same position. However, since the initial composition of the venom resulted from crosses between the ISm and ISy lines, the proteins responsible for a high intensity of a band were likely to be the most abundant proteins in the corresponding protein band of ISm or ISy venom. We therefore used the number of peptides matches from a previous mass spectrometry [18] to classify the proteins in the bands as abundant or not (Table 2). We could identify at least one abundant protein for 12 out of the 17 bands under selection, which are most likely responsible for the observed changes in the overall band intensity.

**Table 2.**
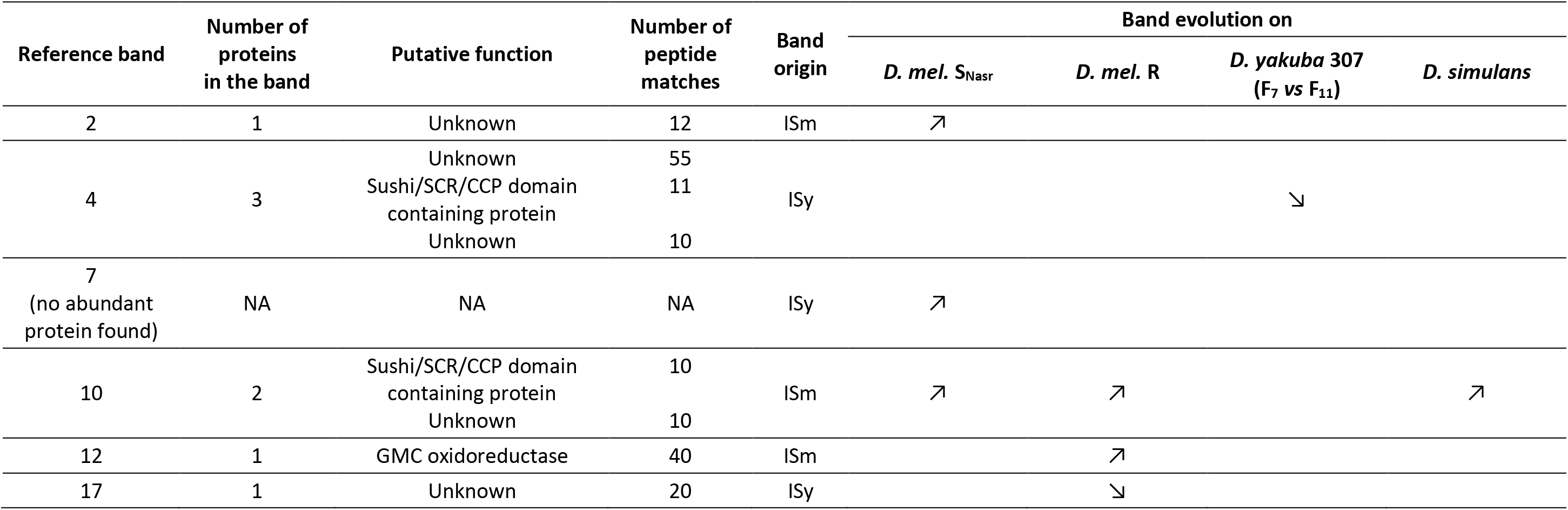
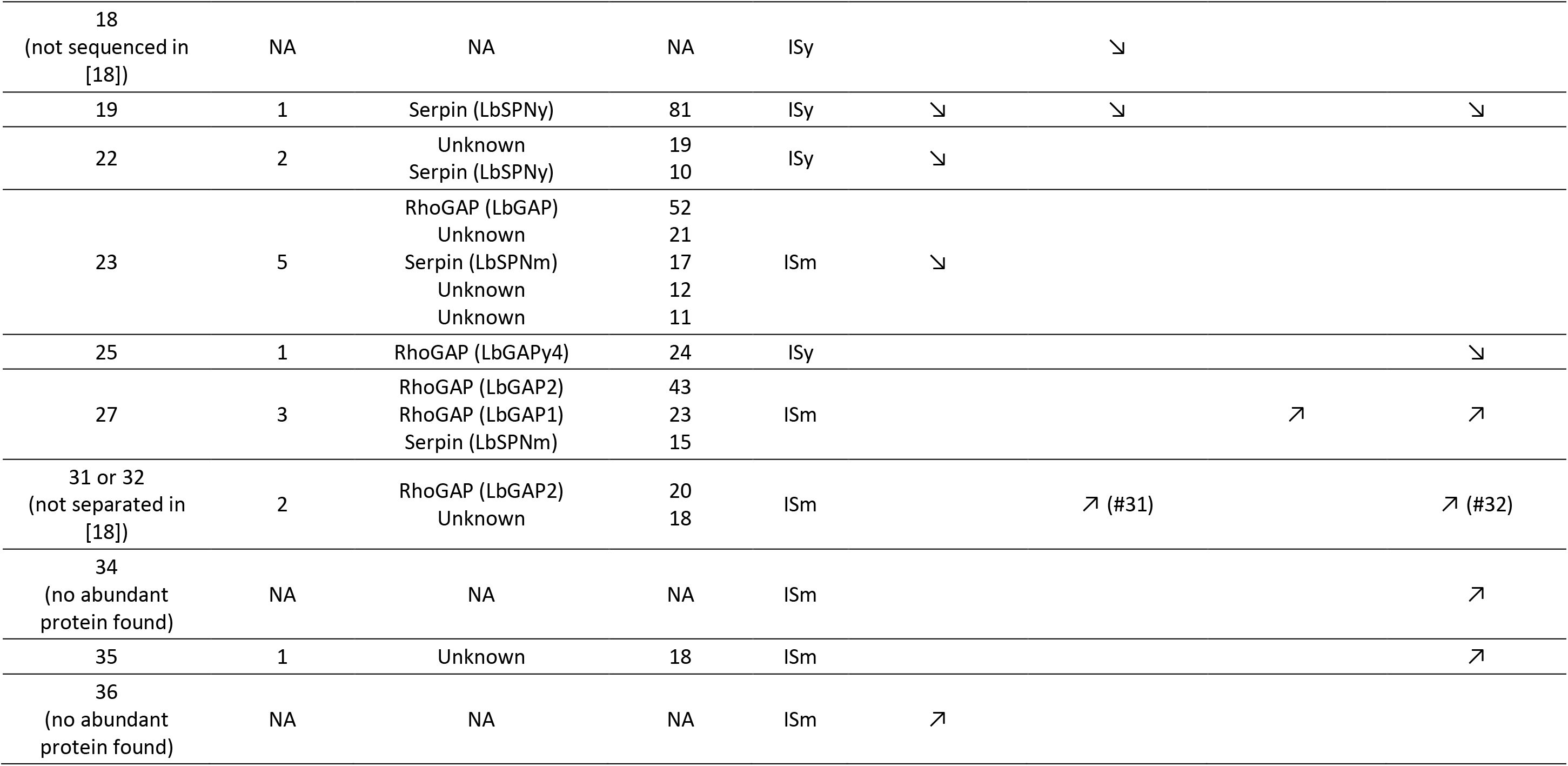
Correspondence between evolving bands and their putative protein content determined from the comparison with data from Colinet et al. (2013a). Only proteins for which at least 10 peptide matches were found in Mascot searches to unisequences identified in transcriptomics of the venom apparatus [18] were considered as abundant and therefore listed. The number of proteins found in the band, their predicted function and the number of peptide matches for each unisequence are provided. Data on the band origin and direction of evolution are from Table 1. Up (“↗”) and down (“↘”) arrows indicate a selection or a counter-selection, respectively. *D. mel.* SNasr: *D. melanogaster* SNasr; *D. mel.* R: *D. melanogaster* R. NA: data not available. On *D. yakuba* 307, the evolution reflects changes between F7 and F11 only.

Although their coding sequence has been previously determined[18], the most abundant proteins in five out of these 12 bands had no similarity to known proteins (Table 2). Their biochemical function thus remains to be determined as for a majority of the proteins contained in the ISm and ISy venoms[18]. Among the bands that evolved on one host only, a Glucose-Methanol-Choline (GMC) oxidoreductase was the most abundant protein in band #12 selected on *D. melanogaster* R (Table 2), and a RhoGAP, LbGAP, was the most abundant protein in band #23 counter-selected on *D. melanogaster* S_Nasr_. We performed a specific analysis for LbGAP that confirmed the counter-selection on *D. melanogaster* S_Nasr_ but also on *D. melanogaster* R (Figure 5A, GLMM, p = 0.009 for *D. melanogaster* S_Nasr_, p = 0.021 for *D. melanogaster* R). Without a good specific antibody effective against the GMC oxidoreductase, no specific analysis could be performed for this protein. The most abundant protein in band #25, counter-selected on *D. simulans*, was another RhoGAP, LbGAPy4. A third RhoGAP, LbGAP2, was found as the most abundant protein in bands #31, selected on *D. melanogaster* R and #32, selected on *D. simulans*, as well as in band #27, selected on *D. simulans* and *D. yakuba* 307 (Table 2). However, the selection/counter-selection of LbGAP2 was not detected in the specific analysis (Figure 5B, LMM, p = 0.14 for *D. melanogaster* R and *D. simulans*, p = 0.20 for *D. melanogaster* S_Nasr_ and p = 0.44 for *D. yakuba* 307).

**Figure 5.**
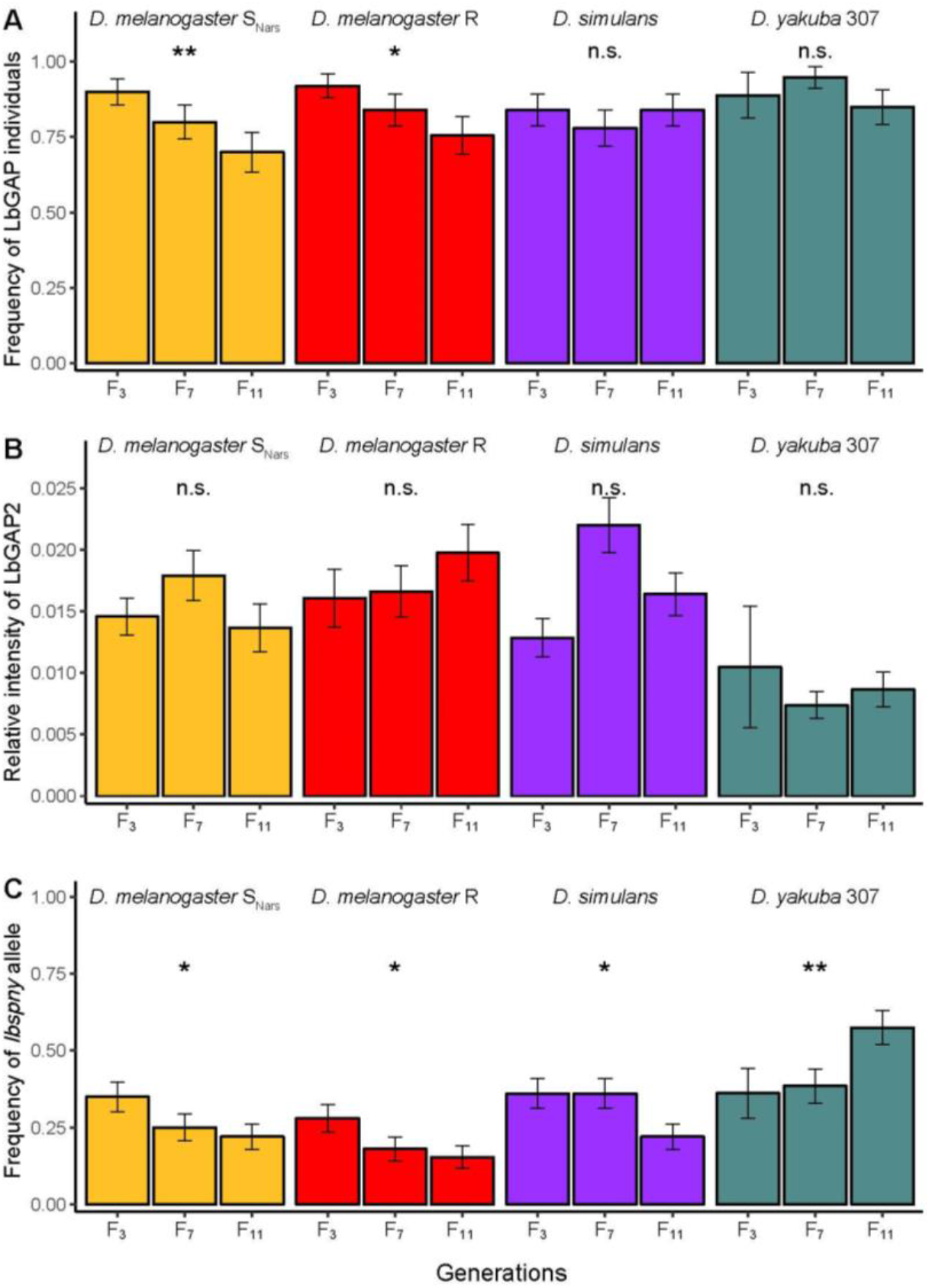
Specific analysis of the evolution of LbGAP, LbGAP2 and LbSPN proteins of parasitoids selected on *D. melanogaster* SNasr (in yellow), *D. melanogaster* R (in red), *D. simulans* (in purple) and *D. yakuba* 307 (in grey-green). A. Evolution of the frequency of individuals harbouring a high quantity of the LbGAP protein. B. Evolution of the LbGAP2 protein amount, relative to the total amount of proteins in the venom samples. C. Evolution of the frequency of the *lbspny* allele.

Regarding the two other bands that evolved on more than one host, a Sushi/SCR/CCP domain-containing protein (SCR: Short Consensus Repeat; CCP: Complement Control Protein) was the most abundant protein in band #10, selected on *D. melanogaster* S_Nasr_, *D. melanogaster* R and *D. simulans* (Table 2). Finally, the most abundant protein in band #19, counter-selected on all hosts except *D. yakuba* 307, was the serpin LbSPNy. The counter-selection of the *lbspny* allele encoding LbSPNy was confirmed by the specific analysis on*D. melanogaster* S_Nasr_, R and *D. simulans* (Figure 5C, GLMM, p = 0.022 for *D. melanogaster* S_Nasr_, p = 0.018 for *D. melanogaster* R, p = 0.015 for *D. simulans*). It also revealed a selection of *lbspny* on *D. yakuba* 307 (Figure 5C, GLMM, p = 0.005) although not detected by the global approach.

### 3.7. Trends of venom evolution

The average venom composition of each of the 44 experimental populations was scaled between 0 and 1 according to their distance to the venom composition of ISm and ISy (Figure 6). When comparing the venom composition in F_3_ and F_11_ generations, we observed a trend for evolution towards the venom composition of ISm for all hosts, except for *D. yakuba* 307 for which the evolution of the venom did not change the relative distance to parental strains (Figure 6, paired Wilcoxon test, p = 0.011 for *D. melanogaster* S_Nasr_, p = 0.010 for *D. melanogaster* R, p = 0.045 for *D. simulans*, p = 0.60 for *D. yakuba* 307).

**Figure 6.**
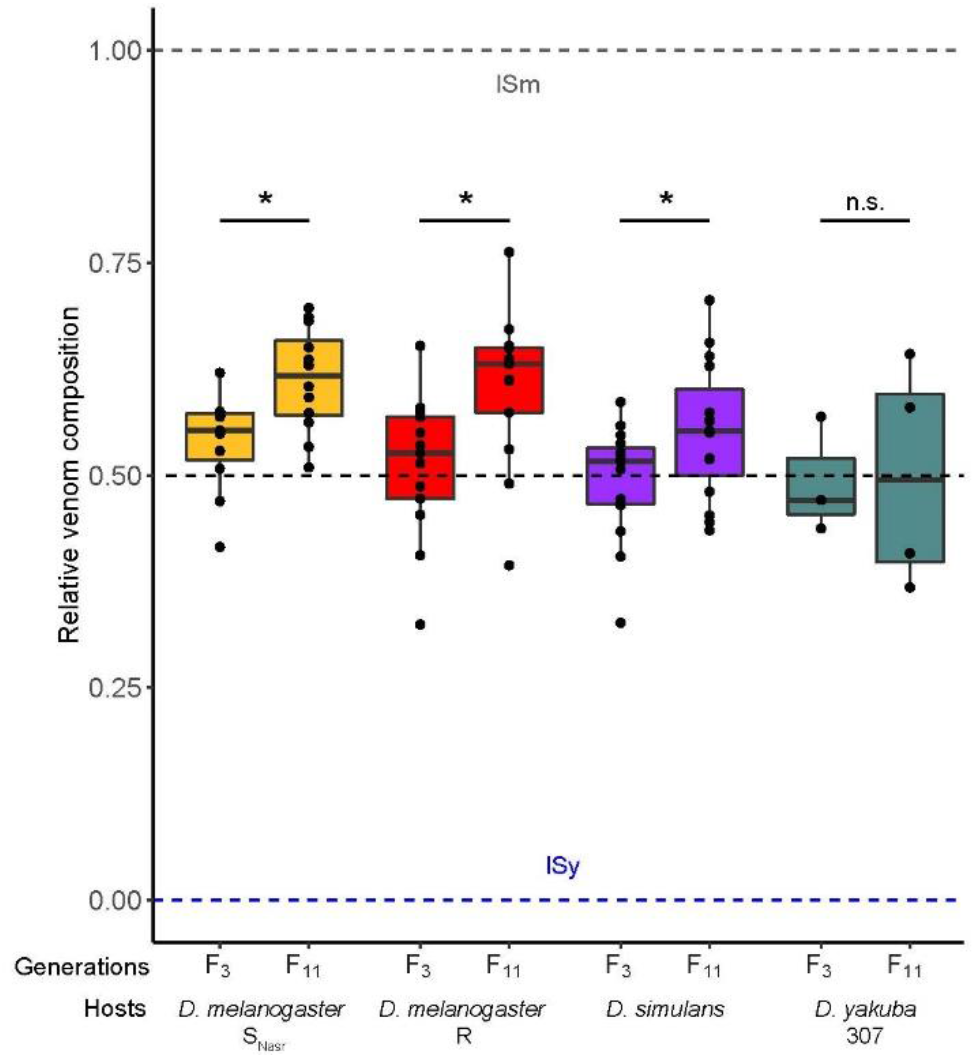
Relative distance between the venom composition of parasitoids selected on *D. melanogaster* SNasr (in yellow), *D. melanogaster* R (in red), *D. simulans* (in purple) and *D. yakuba* 307 (in grey-green) in F3 and F11 and that of *L. boulardi* ISm (1) and ISy (0) lines. The horizontal dashed black line indicates the proportion of ISm alleles after the crossing between ISm and ISy lines (0.5 at the beginning of the experiment). In *D. yakuba* 307, one replicate is missing in F3 because not enough individuals were available to produce the next generation, perform parasitic tests and do the venom analysis.

We then assigned a value between 0 and 1 to each protein band: 0 when a band was present in ISy but absent in ISm, 1 for the opposite (Table 1). In agreement with the above results, most selected protein bands were of the ISm type, whereas most of the counter-selected bands were of the ISy type (Table 2).

## 4. Discussion

This study aimed at testing simultaneously the evolution of (i) the parasitic success and (ii) the venom composition of *L. boulardi* on *Drosophila* hosts from three different species. It was also expected to be informative on whether a venom allowing to develop on different hosts would rather contain broad-spectrum factors or a combination of factors specific to each host. We used an experimental evolution in which selection acted on the standing genetic variation, not on new mutations appeared during the experiment, as is usually done when using fast-growing organisms [45]. The experimental evolution was initiated by crossing *L. boulardi* ISm and ISy parasitoid lines, which differ in their parasitic success on *D. melanogaster* and *D. yakuba* species [8] and their venom composition [18]. Since both lines have spent many generations in the laboratory, the initial variation on which natural selection could act during the experimental evolution resulted mainly from the variation between them. The F_2_ female offspring, whose venom contains new combinations of proteins, were separated in independent populations and reared up to the F_11_ generation on the four different host strains / species on which ISm and/or ISy are successful.

Among these four hosts, *D. melanogaster* S_Nasr_ and *D. simulans* are both susceptible to ISm and ISy. We previously evidenced an evolution of the venom composition of *L. boulardi* not only on a resistant host strain but also on a susceptible one [15]. *D. melanogaster* S_Nasr_ and *D. simulans* were therefore used to determine whether two hosts could exert a different selection pressure on the venom composition of *L. boulardi* despite their susceptibility. Our objective was also i) to confirm that the creation of new combinations of venom factors still allowed parasitic success on these susceptible host strains and ii) to assess whether the venom composition could nevertheless evolve, possibly by selecting still effective but less expensive factors.

A fifth host was used in the experimental evolution, *D. yakuba* 1907 on which neither ISm nor ISy are successful. Interestingly, parasitoids managed to develop on this host until the F_6_ generation suggesting that the creation of new combinations of venom factors at F_2_ has allowed them to develop in a clearly refractory strain for a certain time. However, these new combinations were not successfully selected in any of the replicates since all populations raised on this host ended up being extinct. This may suggest that the virulence factors responsible for this temporary parasitic success are encoded by the same loci leading to a higher fitness of heterozygotes (overdominance), such alleles having been lost due to the small population size.

### 4.1. Parasitic success and venom evolution according to the host

Although the parasitic success remained close to 100% on *D. melanogaster* S_Nasr_ and *D. simulans*, the composition of the parasitoid venom has nevertheless evolved on these susceptible hosts. Although genetic drift due to the small number of individuals per replicate necessarily impacted the venom composition, the observed changes most likely occurred under selection. Genetic drift is a random phenomenon that differently affects each population whereas changes in venom composition were common to most of the replicates suggesting a selection strong enough not to be masked by the drift. The evolution of venom in response to susceptible hosts suggests a selection of some venom components over others, potentially less costly to produce. The venom is not only important to overcome host immune defenses but also to ensure the quality of offspring development [46]. The differential venom evolution on the two susceptible hosts could therefore also result from a selection to increase the fitness of the developing offspring that are facing different susceptible host strains. This scenario would suggest a fine tuning of the host physiology to ensure the best match with the parasitoid larvae requirements.

In contrast, the success of parasitoids selected on *D. melanogaster* R and *D. yakuba* 307 significantly increased with the generation. This rapid increase was not surprising since the selection pressures on parasitoids are very high and it has previously been documented for other host – parasitoid interactions. Accordingly, the parasitic success of *Asobara tabida* increased after seven generations of selection on *D. melanogaster* [47] and that of two aphid parasitoids increased in response to the symbiont-associated resistance of the host [2,3]. A surprising result was the much higher increase in parasitic success for the parasitoids reared on *D. yakuba* 307 than on *D. melanogaster* R, probably because the former was much less successful at F_3_ (about 20% versus 80%). This difference in parasitic success at F_3_ is quite surprising since the ISm line always succeeds on *D. melanogaster* R and ISy on *D. yakuba* 307. It was however consistent with the high extinction rate of F_3_ replicates on *D. yakuba* 307, suggesting a lower virulence of F_2_ parasitoids on this host than on the others. Accordingly, [5,24] identified the virulence of F_1_ (ISm x ISy) hybrids as semi-dominant on *D. melanogaster* and recessive on *D. yakuba*, although they used different strains of *D. melanogaster* and *D. yakuba* than ours.

### 4.2. Specialization of parasitoids to their selection host and differential venom evolution

The greater parasitic success on resistant hosts (*D. yakuba* and *D. melanogaster* R) proved to be specific to the selection host (Figure 2C). Parasitoids maintained on a given resistant host were more successful on this host than those reared on another host. The differential evolution of both parasitic success and venom evolution suggests a specialization of parasitoids on their selection host, as previously reported experimentally in relation to host resistance [3,4] and predicted by simulation results [48]. Such a differential evolution of venom has also been observed in a vertebrate model – the snake of genus *Echis* – in response to diet [49]. The selection of certain protein bands for each host, suggests a role in parasitic success, while the counter-selection suggests a cost associated with their production or function on a given host. These differential selection and counter-selection of the venom content could explain the lesser success of a parasitoid on a resistant host (such as *D. melanogaster* R or *D. yakuba* 307) after a selection on another host strain. For example, the counter-selection of the protein band #19 in the venom of parasitoids selected on *D. melanogaster* S_Nasr_, R, and *D. simulans* may have reduced their success on *D. yakuba* 307. Consistently, another study with other host strains showed that F_1_ (ISm x ISy) hybrids experienced a decrease in virulence against *D. yakuba* after being reared on *D. melanogaster*, suggesting that virulence factors selected on *D. melanogaster* were costly for parasitism on *D. yakuba* [5,50].

A possible hypothesis to explain the differential evolution of venom between parasitoids selected on the two susceptible hosts, *D. melanogaster* S_Nasr_ and *D. simulans* is the occurrence of a host-specific cost to produce certain venom components, resulting in the counter-selection of different proteins. Interestingly, the success on *D. yakuba* of parasitoids selected on *D. melanogaster* S_Nasr_ is lower than those selected on *D. simulans*, confirming the selection of specific venom components to bypass host defenses. It also suggests a difference in immune defenses between each host. As an example, the prophenol-oxidase (PPO) sequences, essential proteins for the melanization process, differ between host species and might be targeted by different venom proteins [51–53].

### 4.3. A combination of mostly host-specific proteins and a few broad-spectrum proteins in venom to succeed on several hosts

The change in intensity observed for most of the 17 venom protein bands which evolved occurred in response to selection by a single host only. This is in accordance with the study of [4] in which most of the differentially expressed genes of selected parasitoids were lineage-specific. As a counter-example, two protein bands evolved in the same direction for *D. melanogaster* S_Nasr_ and R, and *D. simulans* and one for *D. simulans* and *D. yakuba* 307. Therefore, although the number of evolving protein bands may have been underestimated due to a lack of power of the global approach or because they contain several proteins of opposite evolution, our data suggest that the venom of the selected parasitoids contain a combination of mostly host-specific proteins and a few broad-spectrum proteins to succeed on these hosts.

### 4.4. The venom of parasitoids selected on *D. yakuba* 307 evolved more specifically

All PERMANOVAs involving *D. yakuba* 307 showed a significant effect of the “generation × host” interaction suggesting that the venom composition of parasitoids reared on this host evolved more differently than that of those reared on the other hosts. Moreover, parasitoids selected on *D. melanogaster* strains and *D. simulans* showed a trend for evolution towards the venom composition of ISm between F_3_ and F_11_ unlike those selected on *D. yakuba* 307. Likewise, the protein bands #10 and 19, selected and counter-selected on all hosts except *D. yakuba* 307 are of ISm and ISy origin, respectively. In addition, the specific analysis revealed a selection of the *lbspny* allele on *D. yakuba* 307 and a counter-selection on all other hosts. Since *lbspnm* and *lbspny* are two alleles of a co-dominant marker, LbSPN, this could also be interpreted as a selection of *lbspnm* on *D. melanogaster* and *D. simulans* although its role in parasitic success is yet to be demonstrated. These differential trends in the venom composition towards ISm and ISy types may reflect the geographic distribution of parasitoids and hosts - the ISm type of *L. boulardi* being Mediterranean and therefore not encountering *D. yakuba*, mainly present in tropical regions of Africa [5,16,50], as well as the host preferences of the *L. boulardi* lines. ISy was previously shown to oviposit preferentially in *D. yakuba* while ISm preferred *D. melanogaster* [54].

### 4.5. Proteins whose quantity evolved

For many protein bands identified as evolving, the most abundant protein – which is likely responsible for changes in the band intensity – had no predicted function although its coding sequence has been previously determined [18]. These proteins have thus been little or no studied so far but could nevertheless play an essential role in parasitic success and would deserve more attention. Our method without *a priori* seems therefore relevant to identify new candidate proteins, possibly involved in the parasitic success.

Among the proteins with a predicted function and identified as the most abundant one in their evolving band, we found the RhoGAPs LbGAP and LbGAP2 and the serpin LbSPN for which a Western blot analysis with specific antibodies was also performed. The global approach and the specific analysis agreed on the counter-selection of LbGAP on *D. melanogaster* S_Nasr_. This was also evidenced on the resistant strain of *D. melanogaster* R but with the specific analysis only, probably because of the lower power of the global approach. This was surprising since a selection of LbGAP on *D. melanogaster* R was observed in a previous work [15]. However, in this current study, the proportion of LbGAP individuals at F_3_ on *D. melanogaster* hosts was much higher than expected under Hardy-Weinberg equilibrium (90% vs 75%). We therefore cannot exclude a decrease of the frequency of LbGAP individuals to reach equilibrium at F_11_ instead of a counter-selection. Moreover, the initial crosses were done in both directions, instead of only one in the previous study, leading to more balanced frequencies for the overall ISm / ISy alleles. The selection may therefore have acted differently in the two experiments and other alleles than *lbgap* been selected on *D. melanogaster* R. LbGAP is involved in changes of the morphological shape of *D. melanogaster* R lamellocytes, possibly preventing the encapsulation by the host [17,19–22]. We could hypothesize that LbGAP is involved in a strategy to suppress encapsulation. However, we did not observe the increase in parasitoid ability to suppress encapsulation that would therefore be expected under this assumption, but rather an increase of the parasitoid ability to escape from a capsule (Figure 2B). The venom contains several other RhoGAPs including LbGAP2, all mutated on their catalytic site, which might “somehow” compensate for the reduced LbGAP quantity by acting in parasitism success. The selection of LbGAP2 on *D. melanogaster* R, *D. simulans* and *D. yakuba* 307 based on the global approach was however not supported by the specific analysis. This suggests that other proteins in the LbGAP2-containing bands are responsible of the changes in intensity detected by the global approach.

Interestingly, we evidenced the selection of LbSPNy on *D. yakuba* 307, although with the specific analysis only, probably because of the small number of individuals at F_3_ on that host. This is in line with the demonstrated involvement of LbSPNy in the inhibition of the phenoloxidase cascade activation of this same species of *Drosophila* [23]. The phenoloxidase cascade leads to the melanization of the capsule and the release of cytotoxic radicals to kill the parasitoid [23,51–53]. The melanisation process being involved in the strengthening of the capsule, this selection would be rather consistent with the increase of the escape ability of the formed capsule in parasitoids selected on *D. yakuba* associated with the increase in parasitic success (Figure 2B). The global approach and the specific analysis also showed the counter-selection of LbSPNy on *D. melanogaster* S_Nasr_ and R and *D. simulans* in agreement with previous results for *D. melanogaster* hosts [15]. This suggests a possible cost of either the production or the presence of LbSPNy in their venom.

In conclusion, the parasitoid model is very relevant for an experimental evolutionary approach compared to other models of venomous animals such as scorpions or snakes although promising advances were obtained from studies of venomics and virulence on such models [49,55]. Our results have highlighted a specialization of parasitoids on their selection host, in link with a rapid differential evolution of the composition of the venom according to the host. Most of the evolving venom proteins evolved in response to selection by a single host, suggesting that parasitoids use at least partially different mechanisms to bypass the defenses of different hosts, and therefore that these host species may also implement partly different defense mechanisms. *D. melanogaster* and *D. simulans* might share some of them so that part of the venom proteins to succeed on these host species would be common. *D. yakuba* is more apart, the maintenance of parasitoids on this species resulting in the selection of more specific venom factors. From these data, we end up with no universal answer to the question of the venom content of a “generalist” parasitoid in terms of broad-spectrum or host-specific factors. The venom of ISm may contain a mixture of both, allowing success on *D. melanogaster* and *D. simulans*, species usually found in sympatry, while the question remains more open for ISy venom. Finally, in a more general context, the rapid evolution of the venom highlights the strong capacity of parasitoids to possibly adapt to their environment [56]. It is notably striking that crosses between parasitoids creating new combinations of venomous proteins could help them succeed on initially refractory hosts.

## Data accessibility

Data and scripts are available online: https://doi.org/10.6084/m9.figshare.13139090.v1.

Drosophila strains/species and parasitoid lines are maintained in the laboratory and available upon request.

## Author contributions

F.C., H.M-H, J-L.G., D.C. and M.P. designed and planned the experiments. F.C. carried out the experiments. F.C. and H.M-H. analyzed the data. F.C. and D.C. wrote the first draft. All authors were involved in editing of the final version. All authors approve the final version of the manuscript.

## Acknowledgements

We are grateful to Christian Rebuf for help in insect rearing and to Dr. Dominique Joly (Gif/Yvette) for kindly providing the *D. yakuba* strain 307.14. We also thank the “Plateau de microscopie ISA” and Maya Belghazi (CNRS, INP, Marseille, France) for updating the mass spectrometry analyzes. Fanny Cavigliasso PhD was funded by the French Ministry of Higher Education and Research. The work received support from the University of Nice Sophia Antipolis” (Incentive Scientific Credits) and was supported by the French Government (National Research Agency, ANR) through the “Investments for the Future” programs LABEX SIGNALIFE ANR-11-LABX-0028-01 and IDEX UCA Jedi ANR-15-IDEX-01. We thank the three reviewers for their comments on an earlier version of the manuscript. Version 3 of this preprint has been peer-reviewed and recommended by Peer Community In Evolutionary Biology (https://doi.org/10.24072/pci.evolbiol.100124).

## Supplementary

**Table S1.** Extinction of the replicates according to the selection host for the 75 initial experimental populations. For each replicate, F3 to F11 indicate the generation at which none or not enough parasitoids have emerged, thus corresponding to the extinction of the replicate. The F3 generation corresponds to the first generation at which parasitoids emerged from the selection host. The number of replicates at F11 corresponds to the number of replicates (= experimental populations) that were maintained until the end of the experiment.

| Rearing host | Replicates | | | | | | | | | | | | | | | Number of replicates at $F_{11}$ |
| --- | --- | --- | --- | --- | --- | --- | --- | --- | --- | --- | --- | --- | --- | --- | --- | --- |
|  | 1 | 2 | 3 | 4 | 5 | 6 | 7 | 8 | 9 | 10 | 11 | 12 | 13 | 14 | 15 |  |
| *D. melanogaster*<br>$S_{Nasr}$ | $F_6$ | | | | $F_6$ | | | | $F_{11}^*$ | | | | | | | 12 |
| *D. melanogaster*<br>R | | | | | | $F_3$ | | $F_5$ | | | | | | | | 13 |
| *D. yakuba*<br>307 | | $F_3$ | $F_3$ | $F_3$ | $F_3$ | $F_3$ | $F_{11}$ | $F_6$ | | $F_3$ | $F_4$ | | $F_3$ | | $F_3$ | 4 |
| *D. yakuba*<br>1907 | $F_3$ | $F_3$ | $F_3$ | $F_3$ | $F_3$ | $F_3$ | $F_3$ | $F_3$ | $F_6$ | $F_3$ | $F_6$ | $F_3$ | $F_5$ | $F_4$ | $F_3$ | 0 |
| *D. simulans* |  |  |  |  |  |  |  |  |  |  |  |  |  |  |  | 15 |

\*Extinction of this replicate was due to improper handling.

**Table S2.**
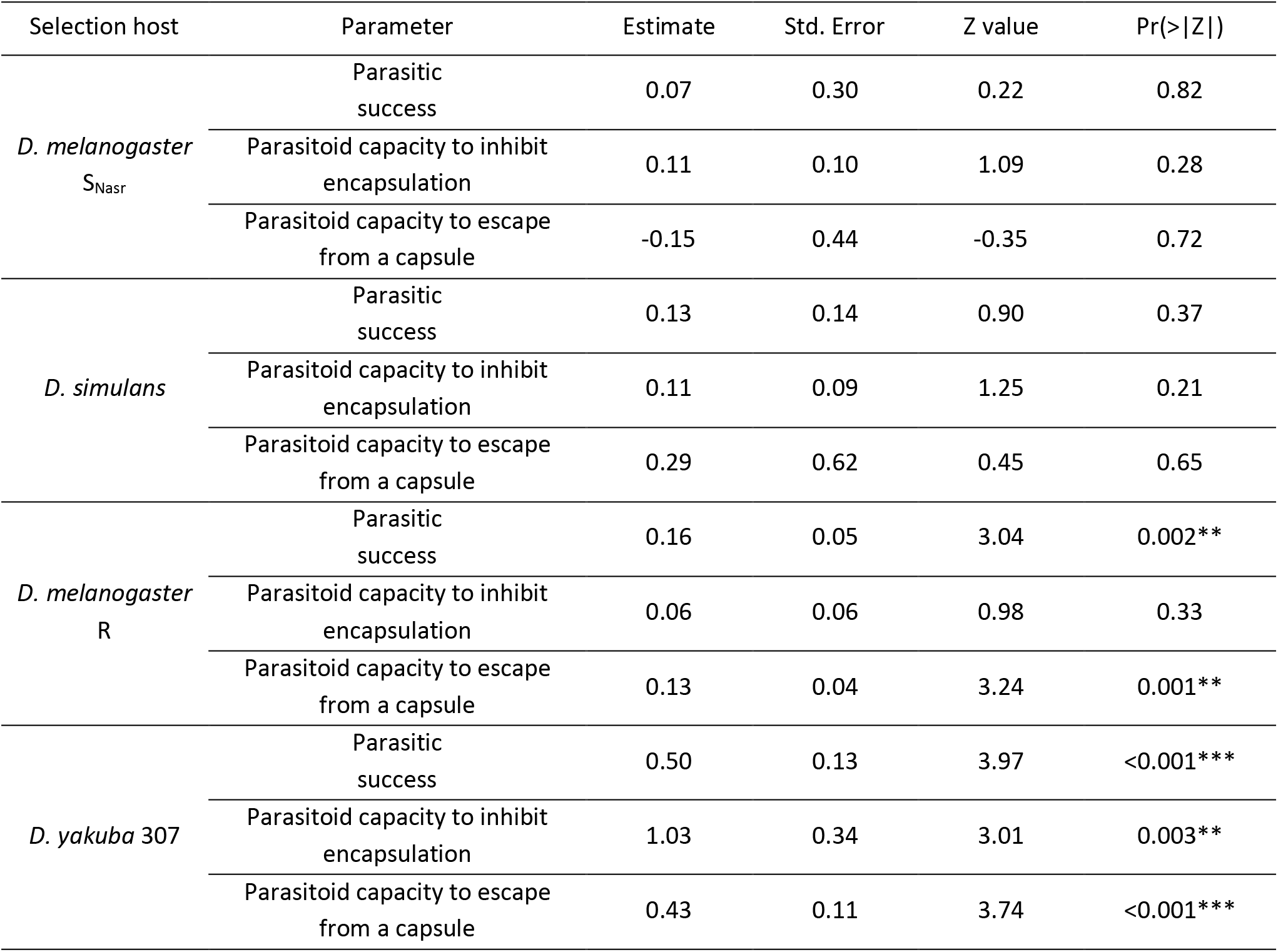
Evolution of the outcome of the interaction according to the selection host. Significance of the generation effect (continuous variable) in GLMM reported in Figure 2B. One model was fitted per host and parameter. Estimate: estimated coefficient, Std. Error: Standard error, Z value: Z statistics, Pr(>|Z|): p-value.

| Selection host | Parameter | Estimate | Std. Error | Z value | $\Pr(>\vert Z\vert )$ |
| --- | --- | --- | --- | --- | --- |
| *D. melanogaster*<br>$S_{Nasr}$ | Parasitic success | 0.07 | 0.30 | 0.22 | 0.82 |
|  | Parasitoid capacity to inhibit encapsulation | 0.11 | 0.10 | 1.09 | 0.28 |
|  | Parasitoid capacity to escape from a capsule | -0.15 | 0.44 | -0.35 | 0.72 |
| *D. simulans* | Parasitic success | 0.13 | 0.14 | 0.90 | 0.37 |
|  | Parasitoid capacity to inhibit encapsulation | 0.11 | 0.09 | 1.25 | 0.21 |
|  | Parasitoid capacity to escape from a capsule | 0.29 | 0.62 | 0.45 | 0.65 |
| *D. melanogaster*<br>R | Parasitic success | 0.16 | 0.05 | 3.04 | 0.002** |
|  | Parasitoid capacity to inhibit encapsulation | 0.06 | 0.06 | 0.98 | 0.33 |
|  | Parasitoid capacity to escape from a capsule | 0.13 | 0.04 | 3.24 | 0.001** |
| *D. yakuba* 307 | Parasitic success | 0.50 | 0.13 | 3.97 | <0.001*** |
|  | Parasitoid capacity to inhibit encapsulation | 1.03 | 0.34 | 3.01 | 0.003** |
|  | Parasitoid capacity to escape from a capsule | 0.43 | 0.11 | 3.74 | <0.001*** |

**Table S3.** Impact of the selection host on the capacity of parasitoids to bypass the defenses of different hosts. Multiple comparisons of means with Tukey post-hoc tests were performed between F11 parasitoids selected on the four hosts for the parasitic success, the parasitoid capacity to inhibit encapsulation and their capacity to escape from a capsule. The capacity to escape from a capsule on *D. melanogaster* SNasr and *D. simulans* was not tested since only three parasitoid females were involved and the statistics were therefore not meaningful. *D. mel.* R or SNasr = *D. melanogaster* R or SNasr; *D. sim*. = *D. simulans*; *D. yak*. = *D. yakuba* 307. Estimate: estimated coefficient, Std. Error: Standard error, Z value: Z statistics, Pr(>|Z|): p-value.

| Tested host | Parameter | Selection host comparison | Estimate | Std. Error | Z value | $Pr(>\vert Z\vert )$ |
| --- | --- | --- | --- | --- | --- | --- |
| *D. melanogaster*<br>$S_{Nasr}$ | Parasitic success | *D. mel. S<sub>Nasr</sub>* – *D. mel. R* | 0.43 | 3.39 | 0.13 | 1.00 |
|  |  | *D. mel. S<sub>Nasr</sub>* – *D. sim.* | 2.25 | 2.99 | 0.75 | 0.87 |
|  |  | *D. mel. S<sub>Nasr</sub>* – *D. yak.* | 2.89 | 3.21 | 0.90 | 0.80 |
|  |  | *D. mel. R* – *D. sim.* | 1.81 | 2.58 | 0.70 | 0.89 |
|  |  | *D. mel. R* – *D. yak.* | 2.45 | 2.59 | 0.95 | 0.77 |
|  |  | *D. sim.* – *D. yak.* | 0.64 | 2.06 | 0.31 | 0.99 |
| *D. melanogaster*<br>$S_{Nasr}$ | Parasitoid capacity to inhibit encapsulation | *D. mel. S<sub>Nasr</sub>* – *D. mel. R* | -1.55 | 2.49 | -0.62 | 0.92 |
|  |  | *D. mel. S<sub>Nasr</sub>* – *D. sim.* | -0.18 | 1.39 | -0.13 | 1.00 |
|  |  | *D. mel. S<sub>Nasr</sub>* – *D. yak.* | 1.80 | 1.37 | 1.31 | 0.54 |
|  |  | *D. mel. R* – *D. sim.* | -1.37 | 2.34 | -0.59 | 0.93 |
|  |  | *D. mel. R* – *D. yak.* | 3.35 | 2.44 | 1.37 | 0.50 |
|  |  | *D. sim.* – *D. yak.* | 1.98 | 1.36 | 1.46 | 0.45 |
| *D. simulans* | Parasitic success | *D. sim.* – *D. mel. S<sub>Nasr</sub>* | 0.92 | 1.50 | 0.61 | 0.92 |
|  |  | *D. sim.* – *D. mel. R* | -1.31 | 2.37 | -0.55 | 0.94 |
|  |  | *D. sim.* – *D. yak.* | -0.83 | 2.64 | -0.31 | 0.99 |
|  |  | *D. mel. S<sub>Nasr</sub>* – *D. mel. R* | -2.23 | 2.56 | -0.87 | 0.81 |
|  |  | *D. mel. S<sub>Nasr</sub>* – *D. yak.* | -1.75 | 2.91 | -0.60 | 0.93 |
|  |  | *D. mel. R* – *D. yak.* | 0.48 | 3.46 | 0.14 | 1.00 |
| *D. simulans* | Parasitoid capacity to inhibit encapsulation | *D. sim.* – *D. mel. S<sub>Nasr</sub>* | 0.10 | 1.23 | 0.08 | 1.00 |
|  |  | *D. sim.* – *D. mel. R* | -1.96 | 2.08 | -0.94 | 0.77 |
|  |  | *D. sim.* – *D. yak.* | 0.70 | 1.35 | 0.52 | 0.95 |
|  |  | *D. mel. S<sub>Nasr</sub>* – *D. mel. R* | -2.06 | 2.26 | -0.91 | 0.79 |
|  |  | *D. mel. S<sub>Nasr</sub>* – *D. yak.* | 0.60 | 1.61 | 0.37 | 0.98 |
|  |  | *D. mel. R* – *D. yak.* | 2.66 | 2.34 | 1.14 | 0.65 |
| *D. melanogaster*<br>R | Parasitic success | *D. mel. R* – *D. sim.* | 1.29 | 0.74 | 1.75 | 0.30 |
|  |  | *D. mel. R* – *D. mel. S<sub>Nasr</sub>* | 1.72 | 0.73 | 2.36 | 0.09 |
|  |  | *D. mel. R* – *D. yak.* | 3.02 | 0.80 | 3.77 | <0.001 *** |
|  |  | *D. mel. S<sub>Nasr</sub>* – *D. sim.* | -0.43 | 0.75 | -0.58 | 0.94 |
|  |  | *D. mel. S<sub>Nasr</sub>* – *D. yak.* | 1.30 | 0.81 | 1.60 | 0.3771 |
|  |  | *D. sim.* – *D. yak.* | 1.73 | 0.80 | 2.17 | 0.13 |

**Table S3.** (continued)
| Tested host | Parameter | Selection host comparison | Estimate | Std. Error | Z value | Pr(>\|Z\|) |
| --- | --- | --- | --- | --- | --- | --- |
| *D. melanogaster*<br>R | Parasitoid capacity to inhibit encapsulation | *D. mel.* R – *D. sim.* | -0.15 | 0.64 | -0.24 | 1.00 |
| | | *D. mel.* R – *D. mel.* $S_{Nasr}$ | 0.22 | 0.65 | 0.34 | 0.99 |
|  |  | *D. mel.* R – *D. yak.* | 2.18 | 0.73 | 2.97 | 0.015 * |
| | | *D. mel.* $S_{Nasr}$ – *D. sim.* | -0.37 | 0.70 | -0.54 | 0.95 |
| | | *D. mel.* $S_{Nasr}$ – *D. yak.* | 1.96 | 0.78 | 2.51 | 0.057 |
|  |  | *D. sim.* – *D. yak.* | 2.33 | 0.77 | 3.02 | 0.013 * |
| *D. melanogaster*<br>R | Parasitoid capacity to escape from a capsule | *D. mel.* R – *D. sim.* | 1.91 | 0.38 | 5.00 | <0.001 *** |
| | | *D. mel.* R – *D. mel.* $S_{Nasr}$ | 1.80 | 0.36 | 5.00 | <0.001 *** |
|  |  | *D. mel.* R – *D. yak.* | 2.12 | 0.42 | 5.04 | <0.001 *** |
| | | *D. mel.* $S_{Nasr}$ – *D. sim.* | 0.12 | 0.38 | 0.31 | 0.99 |
| | | *D. mel.* $S_{Nasr}$ – *D. yak.* | 0.32 | 0.39 | 0.82 | 0.85 |
|  |  | *D. sim.* – *D. yak.* | 0.21 | 0.35 | 0.59 | 0.93 |
| *D. yakuba* 307 | Parasitic success | *D. yak.* – *D. mel.* R | 2.85 | 0.75 | 3.80 | 0.001 ** |
|  |  | *D. yak.* – *D. sim.* | 2.38 | 0.72 | 3.32 | 0.005 ** |
| | | *D. yak.* – *D. mel.* $S_{Nasr}$ | 5.47 | 0.90 | 6.04 | <0.001 *** |
|  |  | *D. mel.* R – *D. sim.* | -0.47 | 0.68 | -0.69 | 0.90 |
| | | *D. mel.* R – *D. mel.* $S_{Nasr}$ | 2.61 | 0.85 | 3.07 | 0.012 * |
| | | *D. sim.* – *D. mel.* $S_{Nasr}$ | 3.08 | 0.83 | 3.70 | 0.001 ** |
| *D. yakuba* 307 | Parasitoid capacity to inhibit encapsulation | *D. yak.* – *D. mel.* R | 0.87 | 1.39 | 0.63 | 0.92 |
|  |  | *D. yak.* – *D. sim.* | -0.19 | 1.13 | -0.17 | 1.00 |
| | | *D. yak.* – *D. mel.* $S_{Nasr}$ | 1.10 | 1.47 | 0.75 | 0.87 |
|  |  | *D. mel.* R – *D. sim.* | -1.07 | 1.03 | -1.04 | 0.72 |
| | | *D. mel.* R – *D. mel.* $S_{Nasr}$ | 0.23 | 1.38 | 0.16 | 1.00 |
| | | *D. sim.* – *D. mel.* $S_{Nasr}$ | 1.29 | 1.05 | 1.23 | 0.60 |
| *D. yakuba* 307 | Parasitoid capacity to escape from a capsule | *D. yak.* – *D. mel.* R | 2.52 | 0.70 | 3.61 | 0.002 ** |
|  |  | *D. yak.* – *D. sim.* | 1.91 | 0.66 | 2.88 | 0.021 * |
| | | *D. yak.* – *D. mel.* $S_{Nasr}$ | 4.90 | 0.85 | 5.80 | <0.001 *** |
|  |  | *D. mel.* R – *D. sim.* | -0.62 | 0.63 | -0.98 | 0.76 |
| | | *D. mel.* R – *D. mel.* $S_{Nasr}$ | 2.38 | 0.80 | 2.99 | 0.015 * |
| | | *D. sim.* – *D. mel.* $S_{Nasr}$ | 3.00 | 0.77 | 3.87 | <0.001 *** |

**Table S4.** Permutational MANOVA for venom evolution on each host strain separately (*D. melanogaster* SNasr, *D. melanogaster* R, *D. yakuba* 307, *D. simulans*). Df: degrees of freedom. Sums of Sqs: sum of squares. F: F statistics. R2: partial R-squared. Pr(>F): p-value based on 5000 constrained permutations within replicates.

| Model | Variance Partition | Df | Sums Of Sqs | F | R<sup>2</sup> | Pr(>F) |
| --- | --- | --- | --- | --- | --- | --- |
| Evolution on<br>*D. melanogaster* $S_{\text{Nasr}}$ host<br>strain | Generation | 1 | 5.70e+09 | 8.57 | 0.044 | 2e-04 *** |
|  | Replicate | 11 | 3.50e+10 | 4.78 | 0.269 | 2e-04 *** |
|  | Residuals | 134 | 8.92e+10 |  | 0.687 |  |
|  | Total | 146<sup>#</sup> | 1.30e+11 |  | 1.000 |  |
| Evolution on<br>*D. melanogaster* R host<br>strain | Generation | 1 | 4.57e+09 | 6.43 | 0.033 | 2e-04 *** |
|  | Replicate | 12 | 3.88e+10 | 4.55 | 0.279 | 2e-04 *** |
|  | Residuals | 135 | 9.60e+10 |  | 0.688 |  |
|  | Total | 148<sup>#</sup> | 1.39e+11 |  | 1.000 |  |
| Evolution on<br>*D. simulans*<br>host strain | Generation | 1 | 2.47e+09 | 3.27 | 0.016 | 0.007 ** |
|  | Replicate | 14 | 5.12e+10 | 4.85 | 0.331 | 2e-04 *** |
|  | Residuals | 134 | 1.01e+11 |  | 0.653 |  |
|  | Total | 149<sup>#</sup> | 1.55e+11 |  | 1.000 |  |
| Evolution on<br>*D. yakuba* 307<br>host strain | Generation | 1 | 1.43e+09 | 1.92 | 0.017 | 0.076 |
|  | Replicate | 3 | 1.88e+10 | 8.44 | 0.228 | 2e-04 *** |
|  | Residuals | 84 | 6.25e+10 |  | 0.755 |  |
|  | Total | 88<sup>#</sup> | 8.28e+10 |  | 1.000 |  |
| Evolution on<br>*D. melanogaster* $S_{\text{Nasr}}$ host<br>strain between F<sub>7</sub> and F<sub>11</sub><br>only | Generation | 1 | 5.70e+09 | 8.57 | 0.044 | 2e-04 *** |
|  | Replicate | 11 | 3.502e+10 | 4.78 | 0.270 | 2e-04 *** |
|  | Residuals | 134 | 8.92e+10 |  | 0.686 |  |
|  | Total | 146 | 1.30e+11 |  | 1.000 |  |
| Evolution on<br>*D. simulans*<br>host strain between F<sub>7</sub><br>and F<sub>11</sub> only | Generation | 1 | 1.39e+09 | 1.99 | 0.013 | 0.074 |
|  | Replicate | 14 | 4.39e+10 | 4.48 | 0.421 | 2e-04 *** |
|  | Residuals | 84 | 5.88e+10 |  | 0.565 |  |
|  | Total | 99 | 1.04e+11 |  | 1.000 |  |
| Evolution on<br>*D. yakuba* 307<br>host strain between F<sub>7</sub><br>and F<sub>11</sub> only | Generation | 1 | 1.61e+09 | 2.30 | 0.022 | 0.042* |
|  | Replicate | 3 | 2.04e+10 | 9.76 | 0.277 | 2e-04 *** |
|  | Residuals | 74 | 5.17e+10 |  | 0.701 |  |
|  | Total | 78<sup>#</sup> | 7.37e+10 |  | 1.000 |  |

<sup>#</sup>: The degrees of freedom vary because different numbers of females were available for venom analysis.

**Table S5.**
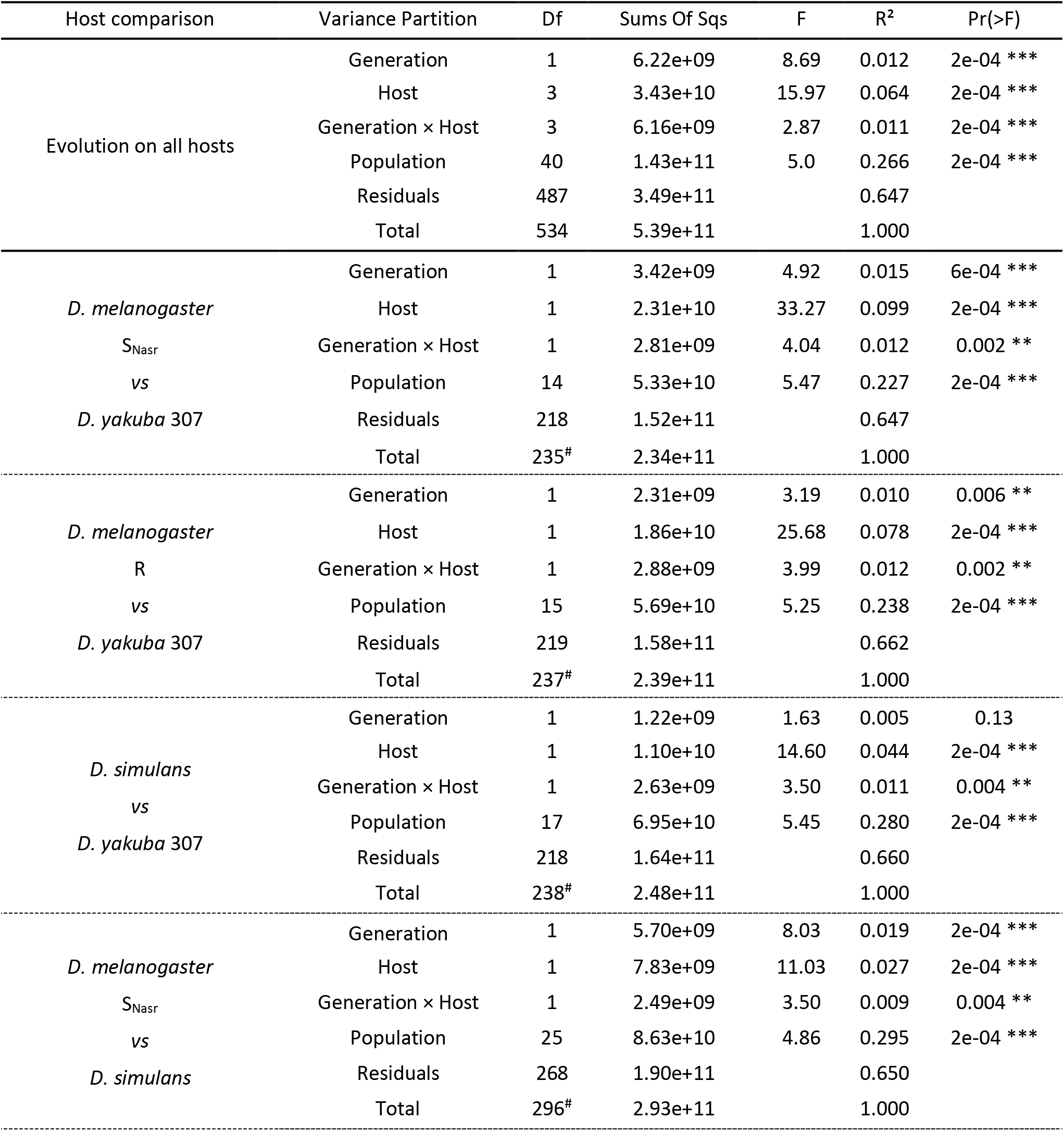
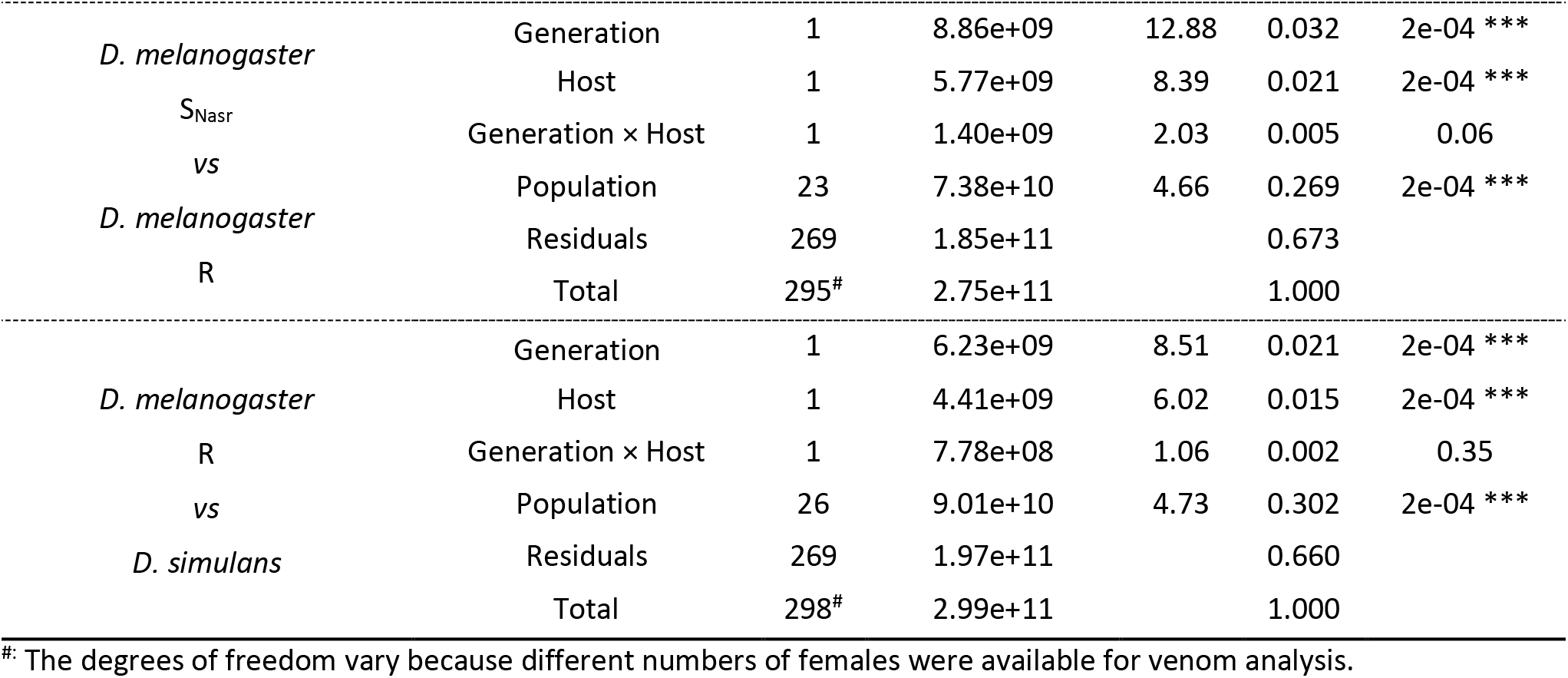
Permutational MANOVA for the evolution of venom on all host strains (*D. melanogaster* SNasr, *D. melanogaster* R, *D. yakuba* 307 and *D. simulans*) and compared 2 by 2. For each effect considered in the model (generation, host strain and the interaction of the two), the following information is provided. Df: degrees of freedom. Sums of Sqs: sum of squares. F: F statistics. R2: partial R-squared. Pr(>F): p-value based on 5000 constrained permutations within replicates. The “population” represents the 44 experimental populations, i.e. replicates maintained on different host strains.

**Tables S6.**
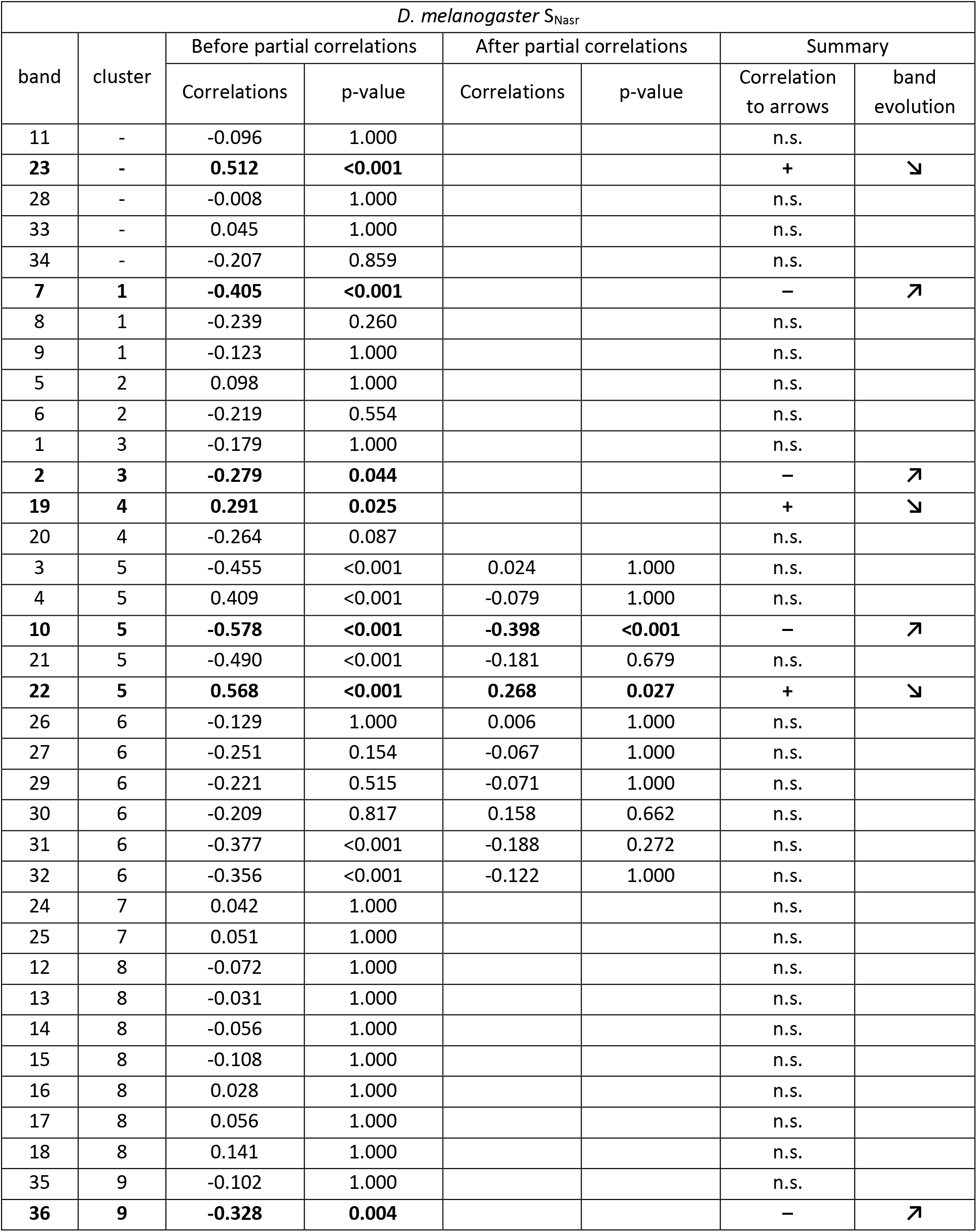
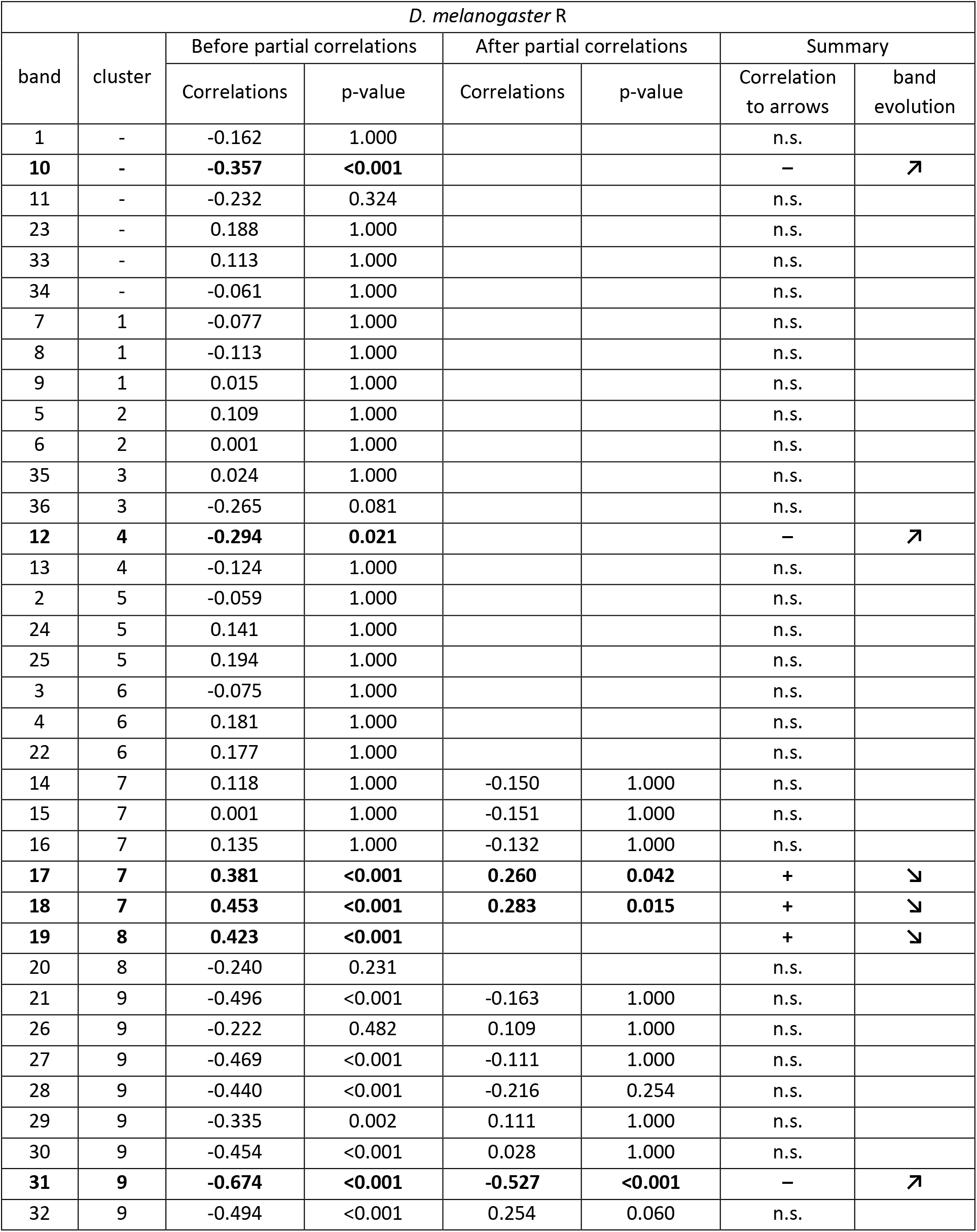
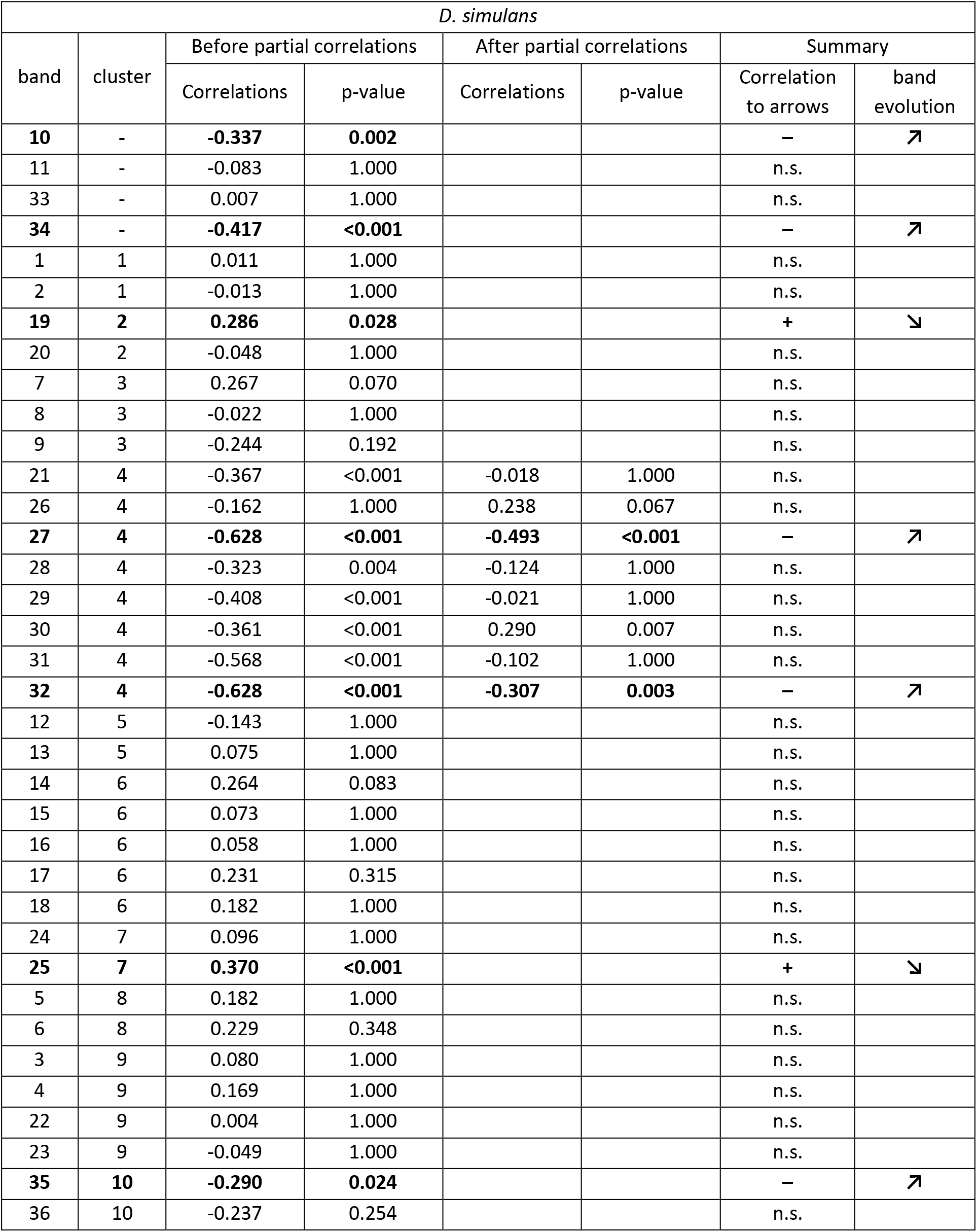
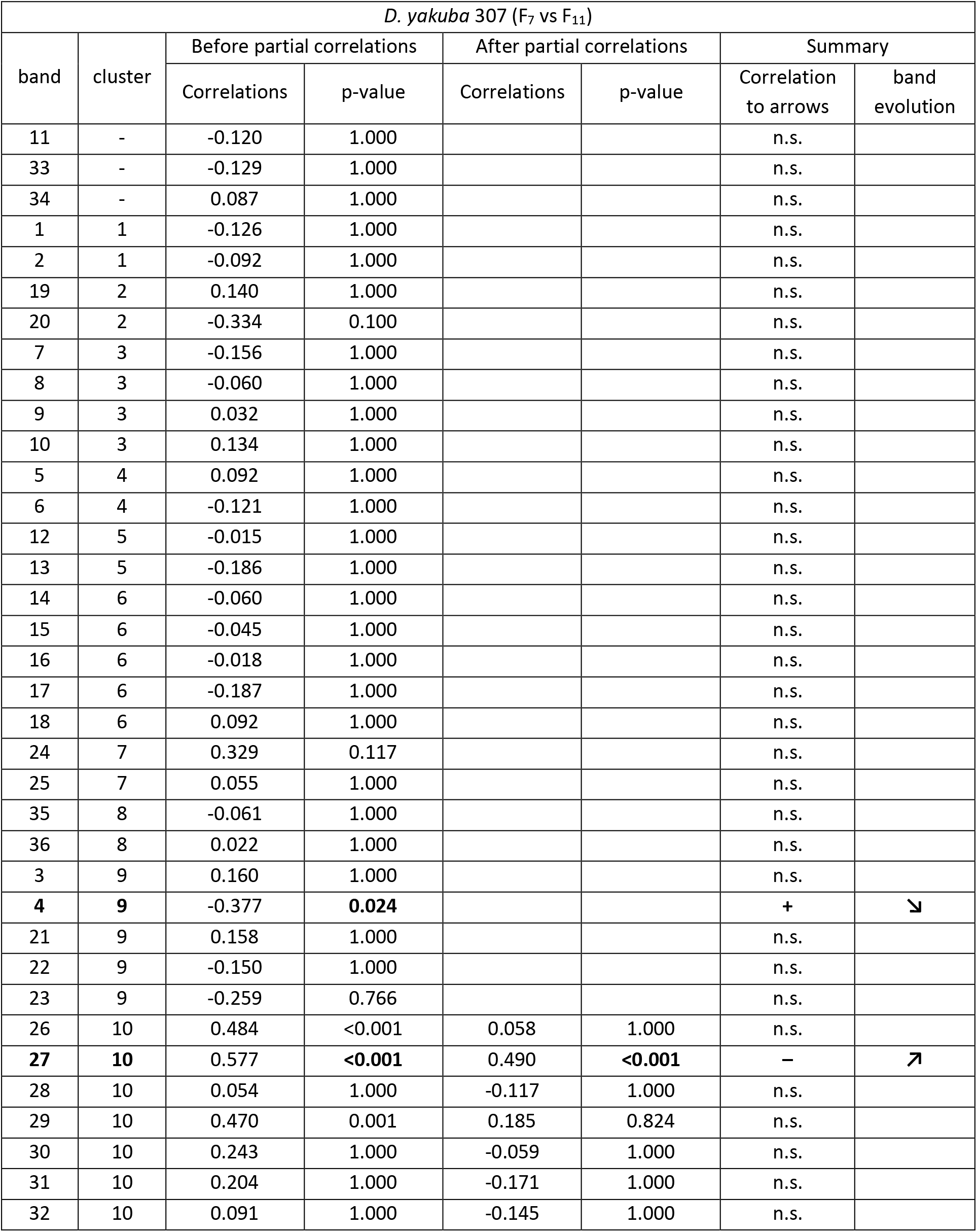
(next four pages) Values of bands correlations of *L. boulardi* parasitoids reared on the four hosts to the arrows from the LDAs for each host separately before and after partial correlation analysis according to the selection host. Cluster numbers come from clustering analysis (see Supplementary Figure S3). n.s.: non-significant. The columns “Correlation to arrows” under “Summary” indicate the significant correlation to the arrows at the end of the analysis. Protein bands in bold represent the evolving protein bands at the end of the analysis. To be considered as an evolving protein band, the sign (+ or –) of the correlation must be the same before and after the partial correlation analysis and the significance level must be under 0.05 before and after the partial correlations. Partial correlations were performed on clusters having at least two protein bands significantly correlated with the arrow. A “↗” in “band evolution” means that the protein band is selected on the corresponding host, a “↘” means that the protein band is counter-selected on the corresponding host. On *D. yakuba* 307, the evolution reflects changes between F7 and F11 only.

| *D. melanogaster* $S_{Nasr}$ | | | | | | | |
| --- | --- | --- | --- | --- | --- | --- | --- |
| band | cluster | Before partial correlations |  | After partial correlations |  | Summary |  |
|  |  | Correlations | p-value | Correlations | p-value | Correlation to arrows | band evolution |
| 11 | - | -0.096 | 1.000 |  |  | n.s. |  |
| **23** | - | **0.512** | **<0.001** |  |  | + | ↘ |
| 28 | - | -0.008 | 1.000 |  |  | n.s. |  |
| 33 | - | 0.045 | 1.000 |  |  | n.s. |  |
| 34 | - | -0.207 | 0.859 |  |  | n.s. |  |
| **7** | **1** | **-0.405** | **<0.001** |  |  | - | ↗ |
| 8 | 1 | -0.239 | 0.260 |  |  | n.s. |  |
| 9 | 1 | -0.123 | 1.000 |  |  | n.s. |  |
| 5 | 2 | 0.098 | 1.000 |  |  | n.s. |  |
| 6 | 2 | -0.219 | 0.554 |  |  | n.s. |  |
| 1 | 3 | -0.179 | 1.000 |  |  | n.s. |  |
| **2** | **3** | **-0.279** | **0.044** |  |  | - | ↗ |
| **19** | **4** | **0.291** | **0.025** |  |  | + | ↘ |
| 20 | 4 | -0.264 | 0.087 |  |  | n.s. |  |
| 3 | 5 | -0.455 | <0.001 | 0.024 | 1.000 | n.s. |  |
| 4 | 5 | 0.409 | <0.001 | -0.079 | 1.000 | n.s. |  |
| **10** | **5** | **-0.578** | **<0.001** | **-0.398** | **<0.001** | - | ↗ |
| 21 | 5 | -0.490 | <0.001 | -0.181 | 0.679 | n.s. |  |
| **22** | **5** | **0.568** | **<0.001** | **0.268** | **0.027** | + | ↘ |
| 26 | 6 | -0.129 | 1.000 | 0.006 | 1.000 | n.s. |  |
| 27 | 6 | -0.251 | 0.154 | -0.067 | 1.000 | n.s. |  |
| 29 | 6 | -0.221 | 0.515 | -0.071 | 1.000 | n.s. |  |
| 30 | 6 | -0.209 | 0.817 | 0.158 | 0.662 | n.s. |  |
| 31 | 6 | -0.377 | <0.001 | -0.188 | 0.272 | n.s. |  |
| 32 | 6 | -0.356 | <0.001 | -0.122 | 1.000 | n.s. |  |
| 24 | 7 | 0.042 | 1.000 |  |  | n.s. |  |
| 25 | 7 | 0.051 | 1.000 |  |  | n.s. |  |
| 12 | 8 | -0.072 | 1.000 |  |  | n.s. |  |
| 13 | 8 | -0.031 | 1.000 |  |  | n.s. |  |
| 14 | 8 | -0.056 | 1.000 |  |  | n.s. |  |
| 15 | 8 | -0.108 | 1.000 |  |  | n.s. |  |
| 16 | 8 | 0.028 | 1.000 |  |  | n.s. |  |
| 17 | 8 | 0.056 | 1.000 |  |  | n.s. |  |
| 18 | 8 | 0.141 | 1.000 |  |  | n.s. |  |
| 35 | 9 | -0.102 | 1.000 |  |  | n.s. |  |
| **36** | **9** | **-0.328** | **0.004** |  |  | - | ↗ |

| *D. melanogaster* R |  |  |  |  |  |  |  |
| --- | --- | --- | --- | --- | --- | --- | --- |
| band | cluster | Before partial correlations |  | After partial correlations |  | Summary |  |
|  |  | Correlations | p-value | Correlations | p-value | Correlation to arrows | band evolution |
| 1 | - | -0.162 | 1.000 |  |  | n.s. |  |
| **10** | - | **-0.357** | **<0.001** |  |  | — | ↗ |
| 11 | - | -0.232 | 0.324 |  |  | n.s. |  |
| 23 | - | 0.188 | 1.000 |  |  | n.s. |  |
| 33 | - | 0.113 | 1.000 |  |  | n.s. |  |
| 34 | - | -0.061 | 1.000 |  |  | n.s. |  |
| 7 | 1 | -0.077 | 1.000 |  |  | n.s. |  |
| 8 | 1 | -0.113 | 1.000 |  |  | n.s. |  |
| 9 | 1 | 0.015 | 1.000 |  |  | n.s. |  |
| 5 | 2 | 0.109 | 1.000 |  |  | n.s. |  |
| 6 | 2 | 0.001 | 1.000 |  |  | n.s. |  |
| 35 | 3 | 0.024 | 1.000 |  |  | n.s. |  |
| 36 | 3 | -0.265 | 0.081 |  |  | n.s. |  |
| **12** | **4** | **-0.294** | **0.021** |  |  | — | ↗ |
| 13 | 4 | -0.124 | 1.000 |  |  | n.s. |  |
| 2 | 5 | -0.059 | 1.000 |  |  | n.s. |  |
| 24 | 5 | 0.141 | 1.000 |  |  | n.s. |  |
| 25 | 5 | 0.194 | 1.000 |  |  | n.s. |  |
| 3 | 6 | -0.075 | 1.000 |  |  | n.s. |  |
| 4 | 6 | 0.181 | 1.000 |  |  | n.s. |  |
| 22 | 6 | 0.177 | 1.000 |  |  | n.s. |  |
| 14 | 7 | 0.118 | 1.000 | -0.150 | 1.000 | n.s. |  |
| 15 | 7 | 0.001 | 1.000 | -0.151 | 1.000 | n.s. |  |
| 16 | 7 | 0.135 | 1.000 | -0.132 | 1.000 | n.s. |  |
| **17** | **7** | **0.381** | **<0.001** | **0.260** | **0.042** | + | ↘ |
| **18** | **7** | **0.453** | **<0.001** | **0.283** | **0.015** | + | ↘ |
| **19** | **8** | **0.423** | **<0.001** |  |  | + | ↘ |
| 20 | 8 | -0.240 | 0.231 |  |  | n.s. |  |
| 21 | 9 | -0.496 | <0.001 | -0.163 | 1.000 | n.s. |  |
| 26 | 9 | -0.222 | 0.482 | 0.109 | 1.000 | n.s. |  |
| 27 | 9 | -0.469 | <0.001 | -0.111 | 1.000 | n.s. |  |
| 28 | 9 | -0.440 | <0.001 | -0.216 | 0.254 | n.s. |  |
| 29 | 9 | -0.335 | 0.002 | 0.111 | 1.000 | n.s. |  |
| 30 | 9 | -0.454 | <0.001 | 0.028 | 1.000 | n.s. |  |
| **31** | **9** | **-0.674** | **<0.001** | **-0.527** | **<0.001** | — | ↗ |
| 32 | 9 | -0.494 | <0.001 | 0.254 | 0.060 | n.s. |  |

| *D. simulans* |  |  |  |  |  |  |  |
| --- | --- | --- | --- | --- | --- | --- | --- |
| band | cluster | Before partial correlations |  | After partial correlations |  | Summary |  |
|  |  | Correlations | p-value | Correlations | p-value | Correlation to arrows | band evolution |
| **10** | - | **-0.337** | **0.002** |  |  | – | ↗ |
| 11 | - | -0.083 | 1.000 |  |  | n.s. |  |
| 33 | - | 0.007 | 1.000 |  |  | n.s. |  |
| **34** | - | **-0.417** | **<0.001** |  |  | – | ↗ |
| 1 | 1 | 0.011 | 1.000 |  |  | n.s. |  |
| 2 | 1 | -0.013 | 1.000 |  |  | n.s. |  |
| **19** | **2** | **0.286** | **0.028** |  |  | + | ↘ |
| 20 | 2 | -0.048 | 1.000 |  |  | n.s. |  |
| 7 | 3 | 0.267 | 0.070 |  |  | n.s. |  |
| 8 | 3 | -0.022 | 1.000 |  |  | n.s. |  |
| 9 | 3 | -0.244 | 0.192 |  |  | n.s. |  |
| 21 | 4 | -0.367 | <0.001 | -0.018 | 1.000 | n.s. |  |
| 26 | 4 | -0.162 | 1.000 | 0.238 | 0.067 | n.s. |  |
| **27** | **4** | **-0.628** | **<0.001** | **-0.493** | **<0.001** | – | ↗ |
| 28 | 4 | -0.323 | 0.004 | -0.124 | 1.000 | n.s. |  |
| 29 | 4 | -0.408 | <0.001 | -0.021 | 1.000 | n.s. |  |
| 30 | 4 | -0.361 | <0.001 | 0.290 | 0.007 | n.s. |  |
| 31 | 4 | -0.568 | <0.001 | -0.102 | 1.000 | n.s. |  |
| **32** | **4** | **-0.628** | **<0.001** | **-0.307** | **0.003** | – | ↗ |
| 12 | 5 | -0.143 | 1.000 |  |  | n.s. |  |
| 13 | 5 | 0.075 | 1.000 |  |  | n.s. |  |
| 14 | 6 | 0.264 | 0.083 |  |  | n.s. |  |
| 15 | 6 | 0.073 | 1.000 |  |  | n.s. |  |
| 16 | 6 | 0.058 | 1.000 |  |  | n.s. |  |
| 17 | 6 | 0.231 | 0.315 |  |  | n.s. |  |
| 18 | 6 | 0.182 | 1.000 |  |  | n.s. |  |
| 24 | 7 | 0.096 | 1.000 |  |  | n.s. |  |
| **25** | **7** | **0.370** | **<0.001** |  |  | + | ↘ |
| 5 | 8 | 0.182 | 1.000 |  |  | n.s. |  |
| 6 | 8 | 0.229 | 0.348 |  |  | n.s. |  |
| 3 | 9 | 0.080 | 1.000 |  |  | n.s. |  |
| 4 | 9 | 0.169 | 1.000 |  |  | n.s. |  |
| 22 | 9 | 0.004 | 1.000 |  |  | n.s. |  |
| 23 | 9 | -0.049 | 1.000 |  |  | n.s. |  |
| **35** | **10** | **-0.290** | **0.024** |  |  | – | ↗ |
| 36 | 10 | -0.237 | 0.254 |  |  | n.s. |  |

| *D. yakuba* 307 (F<sub>7</sub> vs F<sub>11</sub>) |  |  |  |  |  |  |  |
| --- | --- | --- | --- | --- | --- | --- | --- |
| band | cluster | Before partial correlations |  | After partial correlations |  | Summary |  |
|  |  | Correlations | p-value | Correlations | p-value | Correlation to arrows | band evolution |
| 11 | - | -0.120 | 1.000 |  |  | n.s. |  |
| 33 | - | -0.129 | 1.000 |  |  | n.s. |  |
| 34 | - | 0.087 | 1.000 |  |  | n.s. |  |
| 1 | 1 | -0.126 | 1.000 |  |  | n.s. |  |
| 2 | 1 | -0.092 | 1.000 |  |  | n.s. |  |
| 19 | 2 | 0.140 | 1.000 |  |  | n.s. |  |
| 20 | 2 | -0.334 | 0.100 |  |  | n.s. |  |
| 7 | 3 | -0.156 | 1.000 |  |  | n.s. |  |
| 8 | 3 | -0.060 | 1.000 |  |  | n.s. |  |
| 9 | 3 | 0.032 | 1.000 |  |  | n.s. |  |
| 10 | 3 | 0.134 | 1.000 |  |  | n.s. |  |
| 5 | 4 | 0.092 | 1.000 |  |  | n.s. |  |
| 6 | 4 | -0.121 | 1.000 |  |  | n.s. |  |
| 12 | 5 | -0.015 | 1.000 |  |  | n.s. |  |
| 13 | 5 | -0.186 | 1.000 |  |  | n.s. |  |
| 14 | 6 | -0.060 | 1.000 |  |  | n.s. |  |
| 15 | 6 | -0.045 | 1.000 |  |  | n.s. |  |
| 16 | 6 | -0.018 | 1.000 |  |  | n.s. |  |
| 17 | 6 | -0.187 | 1.000 |  |  | n.s. |  |
| 18 | 6 | 0.092 | 1.000 |  |  | n.s. |  |
| 24 | 7 | 0.329 | 0.117 |  |  | n.s. |  |
| 25 | 7 | 0.055 | 1.000 |  |  | n.s. |  |
| 35 | 8 | -0.061 | 1.000 |  |  | n.s. |  |
| 36 | 8 | 0.022 | 1.000 |  |  | n.s. |  |
| 3 | 9 | 0.160 | 1.000 |  |  | n.s. |  |
| **4** | **9** | -0.377 | **0.024** |  |  | **+** | **↘** |
| 21 | 9 | 0.158 | 1.000 |  |  | n.s. |  |
| 22 | 9 | -0.150 | 1.000 |  |  | n.s. |  |
| 23 | 9 | -0.259 | 0.766 |  |  | n.s. |  |
| 26 | 10 | 0.484 | <0.001 | 0.058 | 1.000 | n.s. |  |
| **27** | **10** | 0.577 | **<0.001** | 0.490 | **<0.001** | **-** | **↗** |
| 28 | 10 | 0.054 | 1.000 | -0.117 | 1.000 | n.s. |  |
| 29 | 10 | 0.470 | 0.001 | 0.185 | 0.824 | n.s. |  |
| 30 | 10 | 0.243 | 1.000 | -0.059 | 1.000 | n.s. |  |
| 31 | 10 | 0.204 | 1.000 | -0.171 | 1.000 | n.s. |  |
| 32 | 10 | 0.091 | 1.000 | -0.145 | 1.000 | n.s. |  |

**Figure S1.**
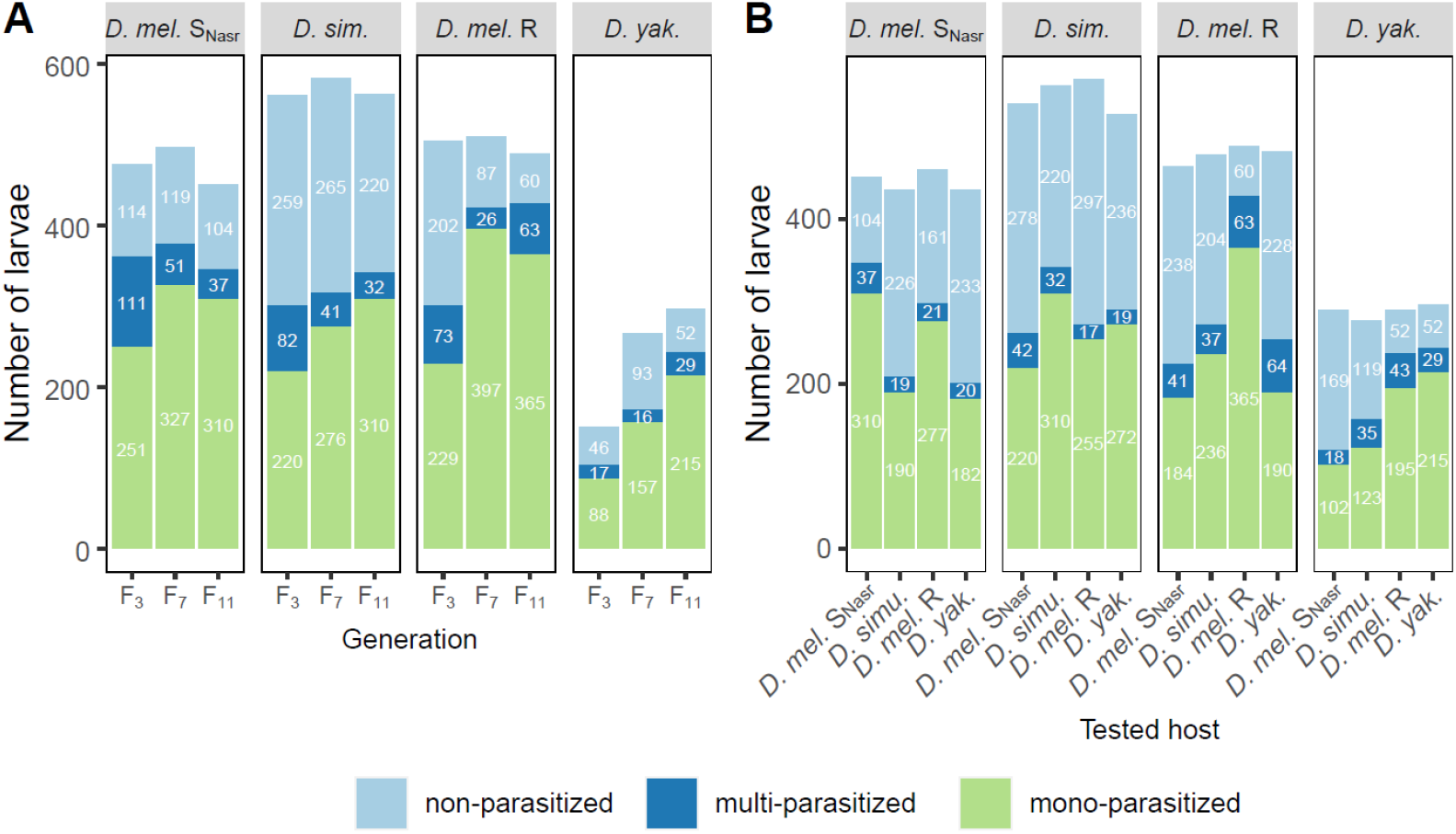
Number and proportion of non-parasitized, multi-parasitized and mono-parasitized larvae dissected for parasitic tests. A. Larvae used to test the evolution of the interaction outcome according to the selection host. B. Larvae used to test for a potential specialization of parasitoids to their selection host.

**Figure S2.**
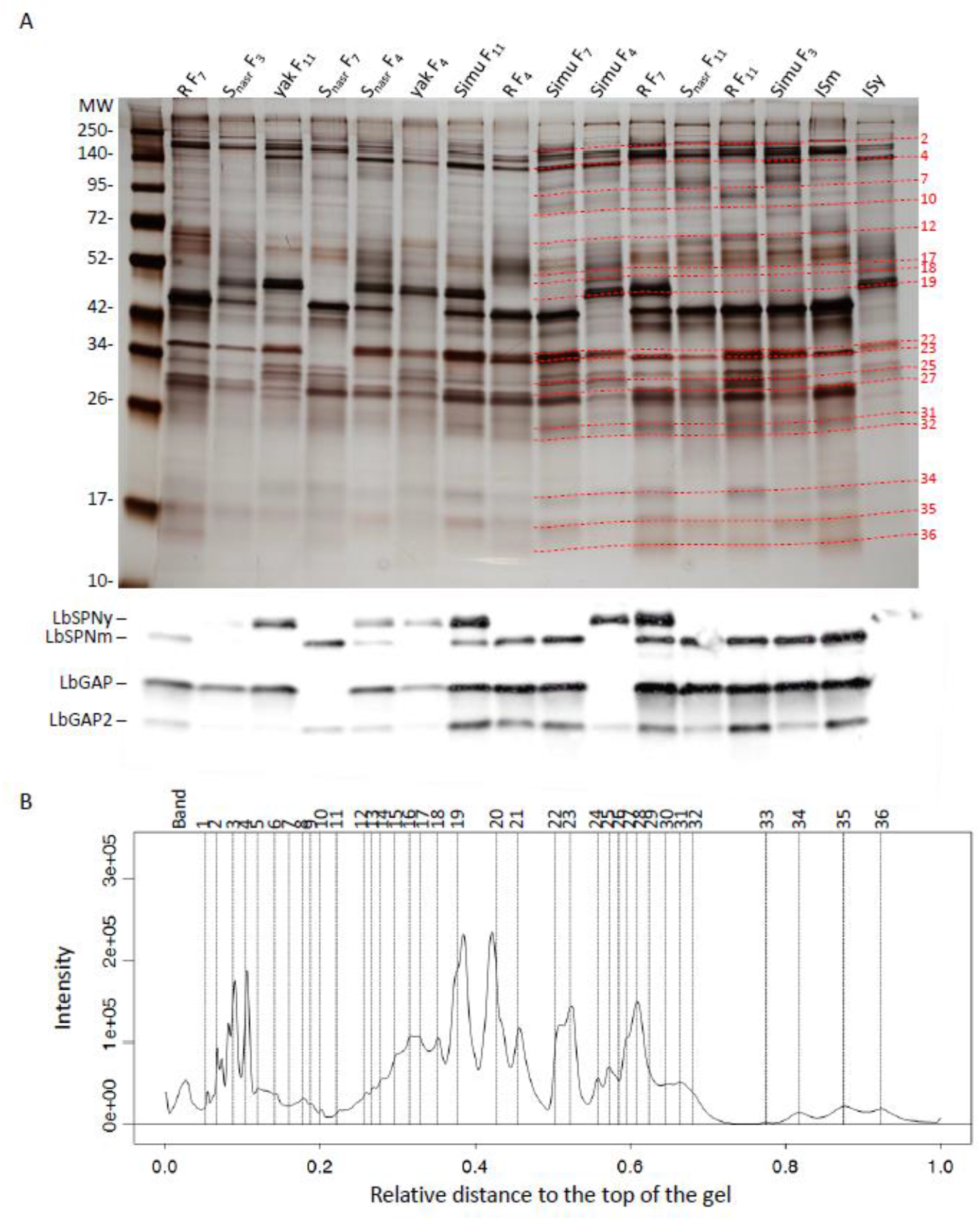
Global analysis of the protein composition of the venom. A. Example of individual venom profiles on silver-stained 1D SDS-PAGE gels and its corresponding western blot with LbSPN, LbGAP and LbGAP2 proteins. Lanes named SNasr, R, yak and Simu correspond to individuals reared on *D. melanogaster* SNasr, *D. melanogaster* R, *D. yakuba* 307, and *D. simulans*, respectively. The numbers in subscript indicate the generation to which the female belongs. ISm and ISy lanes contain the equivalent of half a reservoir but from a pool of sixty gathered reservoirs of ISm and ISy individuals, respectively, used as controls. Red lines (with numbers at right) correspond to the reference bands identified as selected on at least one host. MW: molecular weight in kDa (5 µl, Spectra™ Multicolor Broad Range Protein Ladder, ThermoFisher). B. Simplified mean intensity profile obtained by averaging the intensities of each band over all individual profiles using Phoretix 1D and R functions. Vertical dashed lines correspond to the position of the bands on the gel. Relative distance: distance from the top of the gel relative to the height of the gel. Intensity: intensity of the bands in arbitrary units.

**Figure S3.**
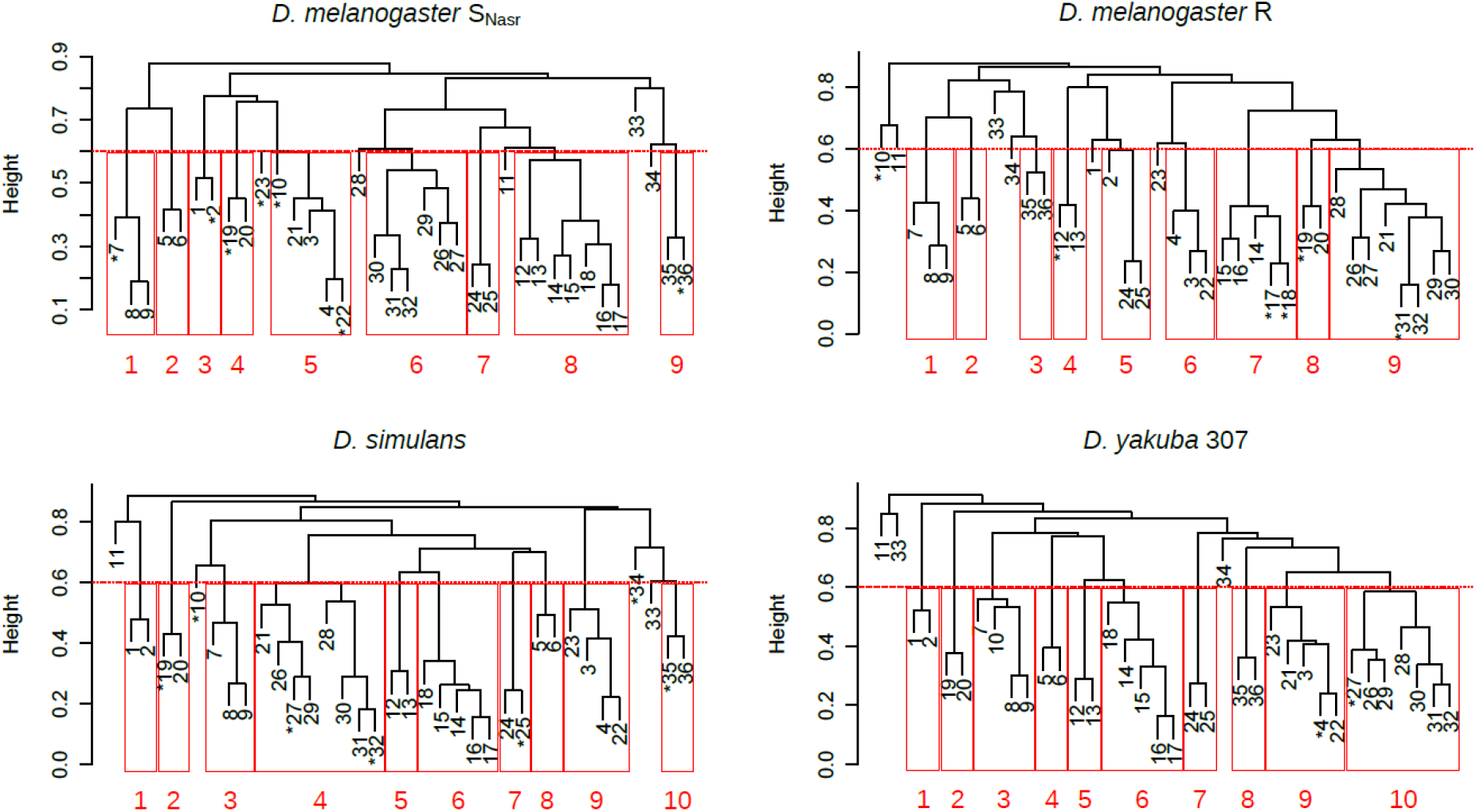
Clustering analysis of protein bands of parasitoids according to their selection host. Each numbered leaf of the dendrogram corresponds to a venom protein band. Bands marked with an asterisk are correlated with the arrow describing the direction of venom evolution after the partial correlation analysis. Height represents the distance between bands calculated as “1 - (absolute value of correlation between bands)”. The horizontal line at 0.6 therefore represents the 0.4 correlation threshold used to build the clusters (in red) for the partial correlation analysis.

**Figure S4.**
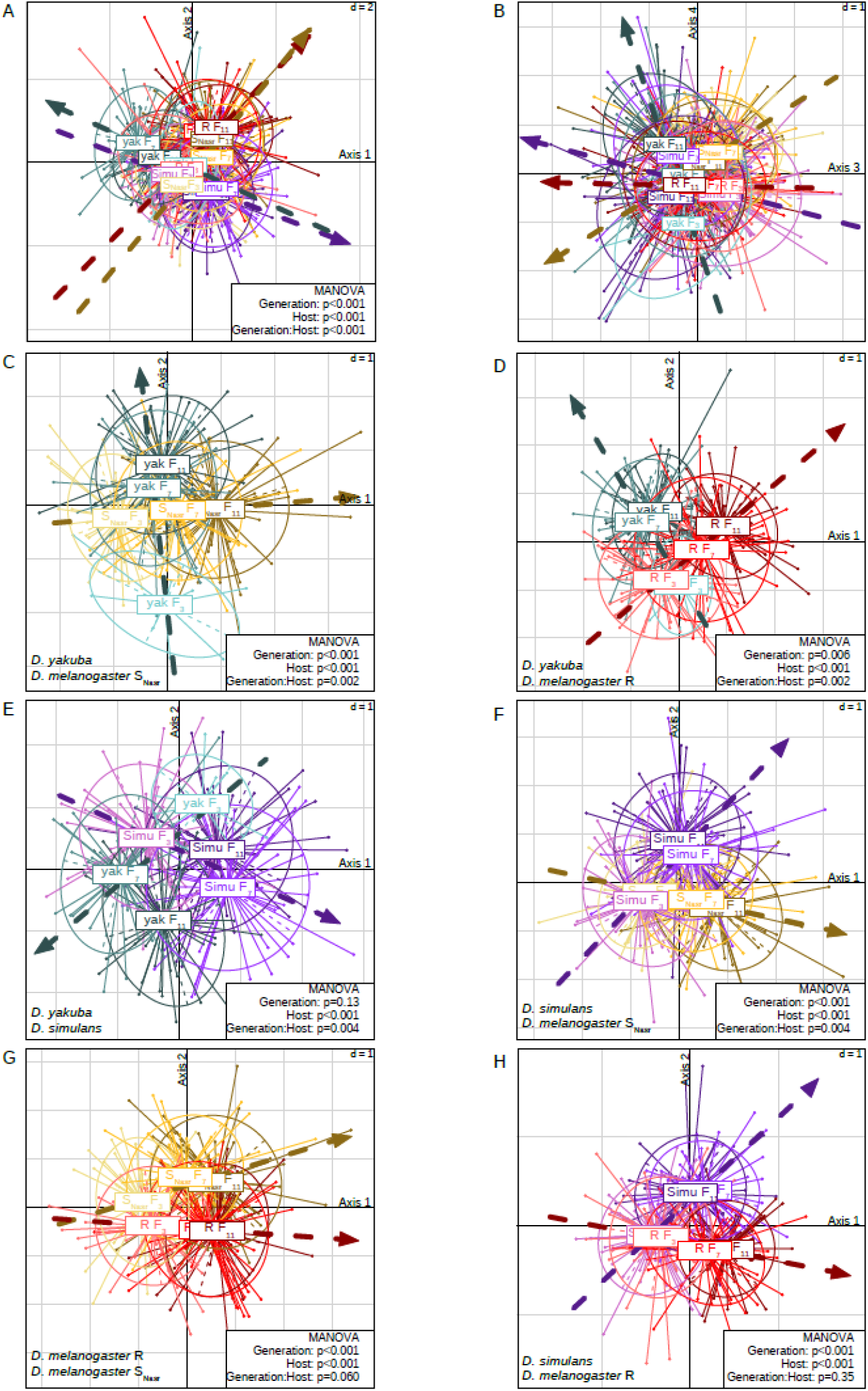
Differential evolution of venom composition. A-B. Position of all the individuals (shown as dots; selected on the four hosts at F3, F7 and F11 generations) on discriminant axes 1 and 2 (A) and discriminant axes 3 and 4 (B). C-H. Position of the individuals on the two first discriminant axes for each two-by-two comparison of venom composition between parasitoids selected on different hosts. For each of these LDAs, the pair of host considered is indicated at the bottom left. Individuals are grouped and coloured according to their selection host and generation. The dotted arrows representing the direction of the venom evolution are the linear regressions fitted to the coordinates of the three centroid points (F3, F7 and F11). P-values obtained from the MANOVA for effects of the “generation”, “host” and “generation × host” interaction are provided at the bottom right of the LDA (see Table S5 for more details). d (top right) represents the scale between two lines.

**Figure S5.**
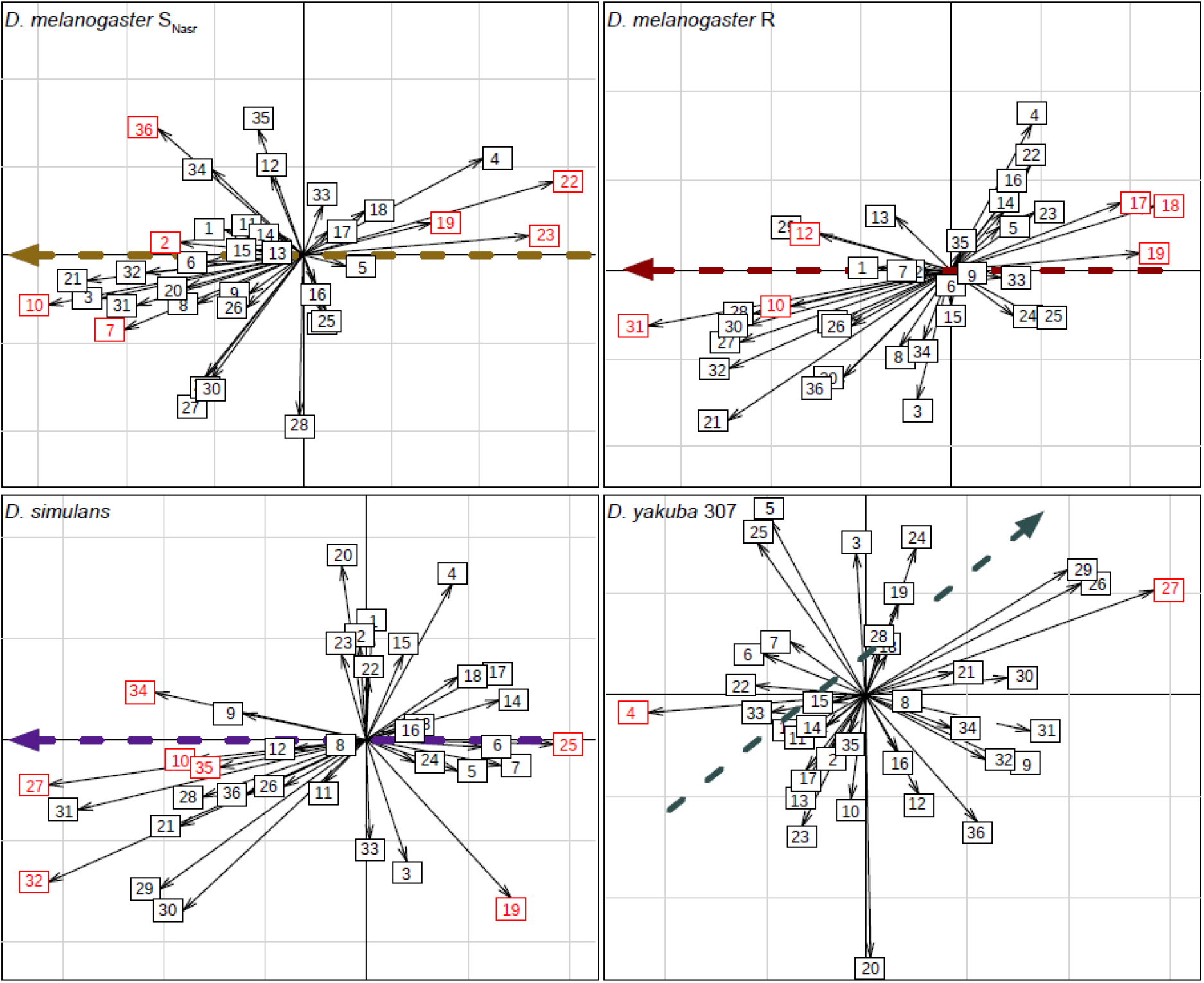
Correlation circles indicating the correlation of protein bands to the linear regressions. The dotted arrows represent the linear regressions calculated from coordinates of the three centroid points corresponding to the F3, F7 and F11 generations and weighted by the number of individuals per generation. For *D. yakuba* 307, the MANOVA revealed a venom evolution only between F7 and F11 so for this host, linear regression was calculated from centroid points of F7 and F11 only. The numbers correspond to the protein bands. A red protein band is a band correlated to the arrow describing the direction of venom evolution on this host, after the partial correlation analysis. Arrows are coloured according to the host. From the center of the axes, the scale between two lines represent a correlation of 0.2.

## References

1. Godfray HCJ. 1994 Parasitoids: behavioral and evolutionary ecology. Princeton Univ. Press, Princeton, NJ.

2. Dion E, Zélé F, Simon J-C, Outreman Y. 2011 Rapid evolution of parasitoids when faced with the symbiont-mediated resistance of their hosts. J. Evol. Biol. 24, 741–750. (doi:10.1111/j.1420-9101.2010.02207.x)

3. Rouchet R, Vorburger C. 2014 Experimental evolution of parasitoid infectivity on symbiont-protected hosts leads to the emergence of genotype specificity. Evolution (N. Y). 68, 1607–1616. (doi:10.1111/evo.12377)

4. Dennis AB, Patel V, Oliver KM, Vorburger C. 2017 Parasitoid gene expression changes after adaptation to symbiont-protected hosts. Evolution (N. Y). 71, 2599–2617. (doi:10.1111/evo.13333)

5. Dupas S, Boscaro M. 1999 Geographic variation and evolution of immunosuppressive genes in a *Drosophila* parasitoid. Ecography (Cop.). 22, 284–291. (doi:10.1111/j.1600-0587.1999.tb00504.x)

6. Fors L, Markus R, Theopold U, Ericson L, Hambäck PA. 2016 Geographic variation and trade-offs in parasitoid virulence. J. Anim. Ecol. 85, 1595–1604. (doi:10.1111/1365-2656.12579)

7. Carton Y, Bouletreau M, Alphen J, Van Lenteren J. 1986 The *Drosophila* parasitic wasps. In The Genetics and Biology of Drosophila, pp. 347–394. Academic Press Inc. (London) Ltd.

8. Dubuffet A, Dupas S, Frey F, Drezen JM, Poirié M, Carton Y. 2007 Genetic interactions between the parasitoid wasp *Leptopilina boulardi* and its *Drosophila* hosts. Heredity (Edinb). 98, 21–27. (doi:10.1038/sj.hdy.6800893)

9. Carton Y, Poirié M, Nappi AJ. 2008 Insect immune resistance to parasitoids. Insect Sci. 15, 67–87. (doi:10.1111/j.1744-7917.2008.00188.x)

10. Nappi AJ. 2010 Cellular immunity and pathogen strategies in combative interactions involving *Drosophila* hosts and their endoparasitic wasps. Invertebr. Surviv. J. 7, 198–210.

11. Vlisidou I, Wood W. 2015 *Drosophila* blood cells and their role in immune responses. FEBS J. 282, 1368–1382. (doi:10.1111/febs.13235)

12. Asgari S, Rivers DB. 2011 Venom proteins from polydnavirus-producing endoparasitoids: Their role in host-parasite interactions. Annu. Rev. Entomol. 56, 313–35. (doi:10.1146/annurev-ento-120709-144849)

13. Poirié M, Colinet D, Gatti JL. 2014 Insights into function and evolution of parasitoid wasp venoms. Curr. Opin. Insect Sci. 6, 52–60. (doi:10.1016/j.cois.2014.10.004)

14. Moreau SJM, Asgari S. 2015 Venom proteins from parasitoid wasps and their biological functions. Toxins (Basel). 7, 2385–2412. (doi:10.3390/toxins7072385)

15. Cavigliasso F, Mathé-Hubert H, Kremmer L, Rebuf C, Gatti J, Malausa T, Colinet D, Poirié M. 2019 Rapid and differential evolution of the venom composition of a parasitoid wasp depending on the host strain. Toxins (Basel). 11, 629.

16. Dubuffet A, Colinet D, Anselme C, Dupas S, Carton Y, Poirié M. 2009 Variation of *Leptopilina boulardi* success in *Drosophila* hosts. What is inside the black box? Adv. Parasitol. 70, 147–188. (doi:10.1016/S0065-308X(09)70006-5)

17. Colinet D, Schmitz A, Cazes D, Gatti JL, Poirié M. 2010 The origin of intraspecific variation of virulence in an eukaryotic immune suppressive parasite. PLoS Pathog. 6, 1–11. (doi:10.1371/journal.ppat.1001206)

18. Colinet D, Deleury E, Anselme C, Cazes D, Poulain J, Azema-Dossat C, Belghazi M, Gatti JL, Poirié M. 2013 Extensive inter- and intraspecific venom variation in closely related parasites targeting the same host: The case of *Leptopilina* parasitoids of *Drosophila*. Insect Biochem. Mol. Biol. 43, 601–611. (doi:10.1016/j.ibmb.2013.03.010)

19. Labrosse C, Stasiak K, Lesobre J, Grangeia A, Huguet E, Drezen JM, Poirié M. 2005 A RhoGAP protein as a main immune suppressive factor in the *Leptopilina boulardi* (Hymenoptera, Figitidae)-*Drosophila melanogaster* interaction. Insect Biochem. Mol. Biol. 35, 93–103. (doi:10.1016/j.ibmb.2004.10.004)

20. Labrosse C, Eslin P, Doury G, Drezen JM, Poirié M. 2005 Haemocyte changes in *D. melanogaster* in response to long gland components of the parasitoid wasp *Leptopilina boulardi*: a Rho-GAP protein as an important factor. J. Insect Physiol. 51, 161–170. (doi:10.1016/j.jinsphys.2004.10.004)

21. Colinet D, Schmitz A, Depoix D, Crochard D, Poirié M. 2007 Convergent use of RhoGAP toxins by eukaryotic parasites and bacterial pathogens. PLoS Pathog. 3, e203. (doi:10.1371/journal.ppat.0030203)

22. Wan B, Goguet E, Ravallec M, Pierre O, Lemauf S, Volkoff A-N, Gatti J-L, Poirié M. 2019 Venom atypical extracellular vesicles as interspecies vehicles of virulence factors involved in host specificity: the case of a *Drosophila* parasitoid wasp. Front. Immunol. 10, 1–14. (doi:10.3389/fimmu.2019.01688)

23. Colinet D, Dubuffet A, Cazes D, Moreau S, Drezen JM, Poirié M. 2009 A serpin from the parasitoid wasp *Leptopilina boulardi* targets the *Drosophila* phenoloxidase cascade. Dev. Comp. Immunol. 33, 681–689. (doi:10.1016/j.dci.2008.11.013)

24. Dupas S, Frey F, Carton Y. 1998 A single parasitoid segregating factor controls immune suppression in *Drosophila*. J. Hered. 89, 306–311. (doi:10.1093/jhered/89.4.306)

25. Carton Y, Frey F, Nappi A. 1992 Genetic determinism of the cellular immune reaction in *Drosophila melanogaster*. Heredity (Edinb). 69, 393–399. (doi:10.1038/hdy.1992.141)

26. Carton Y, Nappi AJ. 1997 *Drosophila* cellular immunity against parasitoids. Parasitol. Today 13, 218– 227.

27. Poirié M, Frey F, Hita M, Huguet E, Lemeunier F, Periquet G, Carton Y. 2000 *Drosophila* resistance genes to parasitoids: Chromosomal location and linkage analysis. Proc. R. Soc. B Biol. Sci. 267, 1417– 1421. (doi:10.1098/rspb.2000.1158)

28. Kim-Jo C, Gatti JL, Poirié M. 2019 *Drosophila* cellular immunity against parasitoid wasps: A complex and time-dependent process. Front. Physiol. 10, 603. (doi:10.3389/fphys.2019.00603)

29. Hita M, Espagne E, Lemeunier F, Pascual L, Carton Y, Periquet G, Poirié M. 2006 Mapping candidate genes for *Drosophila melanogaster* resistance to the parasitoid wasp *Leptopilina boulardi*. Genet. Res. 88, 81–91. (doi:10.1017/S001667230600841X)

30. Fellowes MDE, Kraaijeveld AR, Godfray HCJ. 1998 Trade-off associated with selection for increased ability to resist parasitoid attack in *Drosophila melanogaster*. Proc. R. Soc. B Biol. Sci. 265, 1553–1558. (doi:10.1098/rspb.1998.0471)

31. Singmann H, Bolker B, Westfall J, Aust F. 2019 afex: analysis of factorial experiments.

32. Chung Y, Rabe-Hesketh S, Dorie V, Gelman A, Liu J. 2013 A nondegenerate penalized likelihood estimator for variance parameters in multilevel models. Psychometrika 78, 685–709.

33. Bolker BM, Brooks ME, Clark CJ, Geange SW, Poulsen JR, Stevens MHH, White JSS. 2009 Generalized linear mixed models: a practical guide for ecology and evolution. Trends Ecol. Evol. 24, 127–135. (doi:10.1016/j.tree.2008.10.008)

34. Harrison XA. 2015 A comparison of observation-level random effect and Beta-Binomial models for modelling overdispersion in Binomial data in ecology & evolution. PeerJ 2015. (doi:10.7717/peerj.1114)

35. Hothorn T, Bretz F, Westfall P. 2008 Simultaneous inference in general parametric models. Biometrical J. 50, 346–363. (doi:10.1002/bimj.200810425)

36. Mathé-Hubert H, Gatti JL, Colinet D, Poirié M, Malausa T. 2015 Statistical analysis of the individual variability of 1D protein profiles as a tool in ecology: An application to parasitoid venom. Mol. Ecol. Resour. 15, 1120–1132. (doi:10.1111/1755-0998.12389)

37. Colinet D, Mathé-Hubert H, Allemand R, Gatti JL, Poirié M. 2013 Variability of venom components in immune suppressive parasitoid wasps: From a phylogenetic to a population approach. J. Insect Physiol. 59, 205–212. (doi:10.1016/j.jinsphys.2012.10.013)

38. Colinet D, Kremmer L, Lemauf S, Rebuf C, Gatti JL, Poirié M. 2014 Development of RNAi in a *Drosophila* endoparasitoid wasp and demonstration of its efficiency in impairing venom protein production. J. Insect Physiol. 63, 56–61. (doi:10.1016/j.jinsphys.2014.02.011)

39. Dixon P. 2003 VEGAN, a package of R functions for community ecology. J. Veg. Sci. 14, 927–930.

40. Anderson MJ. 2017 Permutational Multivariate Analysis of Variance (PERMANOVA). Wiley StatsRef Stat. Ref. Online, 1–15. (doi:10.1002/9781118445112.stat07841)

41. Dray S, Dufour A-B. 2007 The ade4 package: implementing the duality diagram for ecologists. J. Stat. Softw. 22, 1–20.

42. Pinheiro J, Bates D, DebRoy S, D S, Team RDC. 2011 nlme: linear and nonlinear mixed effects models. R Packag. version 3, 1–97.

43. Qiao Z, Zhou L, Huang J. 2008 Effective linear discriminant analysis for high dimensional, low sample size data. Proceeding World Congr. Eng. 2, 2–4.

44. Luo D, Ding C, Huang H. 2011 Linear discriminant analysis: New formulations and overfit analysis. Proc. Natl. Conf. Artif. Intell. 1, 417–422.

45. Lenski RE. 2017 Experimental evolution and the dynamics of adaptation and genome evolution in microbial populations. ISME J. 11, 2181–2194. (doi:10.1038/ismej.2017.69)

46. Mrinalini, Siebert AL, Wright J, Martinson E, Wheeler D, Werren JH. 2015 Parasitoid venom induces metabolic cascades in fly hosts. Metabolomics 11, 350–366. (doi:10.1007/s11306-014-0697-z)

47. Kraaijeveld AR, Hutcheson KA, Limentani EC, Godfray HCJ. 2001 Costs of counterdefenses to host resistance in a parasitoid of *Drosophila*. Evolution (N. Y). 55, 1815–1821. (doi:10.1111/j.0014-3820.2001.tb00830.x)

48. Lapchin L. 2002 Host-parasitoid association and diffuse coevolution: When to be a generalist? Am. Nat. 160, 245–254. (doi:10.1086/341020)

49. Barlow A, Pook CE, Harrison RA, Wüster W. 2009 Coevolution of diet and prey-specific venom activity supports the role of selection in snake venom evolution. Proc. R. Soc. B Biol. Sci. 276, 2443–2449. (doi:10.1098/rspb.2009.0048)

50. Dupas S, Dubuffet A, Carton Y, Poirié M. 2009 Local, geographic and phylogenetic scales of coevolution in *Drosophila*-parasitoid interactions. Adv. Parasitol. 70, 281–295. (doi:10.1016/S0065-308X(09)70011-9)

51. Dudzic JP, Kondo S, Ueda R, Bergman CM, Lemaitre B. 2015 *Drosophila* innate immunity: Regional and functional specialization of prophenoloxidases. BMC Biol. 13, 1–16. (doi:10.1186/s12915-015-0193-6)

52. Nappi AJ, Christensen BM. 2005 Melanogenesis and associated cytotoxic reactions: Applications to insect innate immunity. Insect Biochem. Mol. Biol. 35, 443–459. (doi:10.1016/j.ibmb.2005.01.014)

53. Cerenius L, Lee BL, Söderhäll K. 2008 The proPO-system: pros and cons for its role in invertebrate immunity. Trends Immunol. 29, 263–271. (doi:10.1016/j.it.2008.02.009)

54. Dubuffet A, Álvarez CIR, Drezen JM, Van Alphen JJM, Poirié M. 2006 Do parasitoid preferences for different host species match virulence? Physiol. Entomol. 31, 170–177. (doi:10.1111/j.1365-3032.2006.00505.x)

55. Evans ERJ, Northfield TD, Daly NL, Wilson DT. 2019 Venom costs and optimization in scorpions. Front. Ecol. Evol. 7, 1–7. (doi:10.3389/fevo.2019.00196)

56. Mathé-Hubert H, Kremmer L, Colinet D, Gatti JL, Van Baaren J, Delava É, Poirié M. 2019 Variation in the venom of parasitic wasps, drift, or selection? Insights from a multivariate QST analysis. Front. Ecol. Evol. 7, 256. (doi:10.3389/fevo.2019.156)

